# Peracetic Acid Efficacy and Decay Kinetics in Poultry Processing under Chiller Conditions

**DOI:** 10.1101/2025.09.04.674287

**Authors:** Vyshnavi Ciluveru, Jason Simon, Jeffery M. Farber, Shawn D. Ryan, Chandrasekhar R. Kothapalli, Daniel S. Munther

## Abstract

Pathogens on poultry products continue to pose critical public health risks. Despite significant literature examining sanitizer impact on bacterial pathogens during chilling, the mechanisms of sanitizer efficacy in terms of pathogen and organic load, sanitizer levels, and exposure duration are not well understood. To assess these, we report on experimentally-informed-mechanistic-modeling to describe pathogen dynamics during poultry chilling. The shedding and survival of a five-strain cocktail of poultry-plant derived *Salmonella enterica* serovars, at high and low loads, with exposure to peracetic acid (PAA; 0 – 200 mg·L^-1^) for up to 60 min in 10-L chiller tanks, in the presence or absence of whole chicken carcass/parts, were measured. Process water parameters versus time were simultaneously monitored. Results suggest that total dissolved solids (TDS) predict PAA decay more consistently than chemical oxygen demand (COD). A mathematical model for PAA decay and pathogen shedding/inactivation was developed. This model accurately predicted PAA level changes in the pre/main chiller of a high-speed poultry processing plant in North America. Without PAA, *Salmonella* shedding from chicken thighs is influenced by rinse time and number of rinses. Without organic load, residual PAA (1 mg·L^-1^) inactivated bacteria given sufficient exposure time, although PAA levels > 5 mg·L^-1^ were essential for rapid inactivation. With organic load, initial PAA concentration (> 40 mg·L^-1^) and exposure time (> 2 min) were critical for bacterial inactivation, with model results connecting process conditions to dominate modes of bacterial inactivation on chicken. The insights from such experimental-modeling studies provide key tools for processors to improve pathogen control during chilling.

## 1. Introduction

The United States led global chicken meat production in 2023 with approximately 21 million metric tons being produced (USDA, 2023). The per capita consumption of broiler meat in the United States reached approximately 97 lbs in 2022, reflecting a steady increase since 2010 [1, 2]. In the developing nations, growing demand for poultry meat necessitates production on a larger scale. However, without adequate infrastructure to ensure proper disinfection and hygienic handling, large-scale production may lead to an increased prevalence of enteric pathogens in the final products, raising the risk of their transmission from farms to consumers. Ensuring food safety and maintaining disinfection standards are critical to deliver safe poultry products to consumers [3].

A major public health concern is pathogen contamination during poultry processing. For instance, *Salmonella enterica* is continuously shed into the environment via animal feces and can survive in dried fecal material, soil, and vegetation for extended periods of time [4]. *Salmonella* is frequently linked to poultry farms and poultry products, and poses significant risks due to its ability to contaminate, survive, and grow during production, processing, and handling stages, with improper storage and cooking providing conditions for its persistence and spread [5]. The economic burden of poultry-associated salmonellosis in the US is substantial, with annual costs estimated at $2.8 billion, highlighting the critical need for enhanced safety measures across the supply chain [6]. A study estimated that around 10% − 22% of human salmonellosis cases in developed countries are linked to the consumption of contaminated poultry products [7]. In developing countries, *Salmonella* and *Campylobacter* are consistently detected in poultry meat and environmental samples, and frequently identified in small-scale poultry slaughter facilities, processing units, pluck shops, cottage poultry processors, local retail markets, and wet markets [4, 8].

Surveillance reports from the USDA-FSIS, NARMS, and various independent studies indicate that multiple *Salmonella* serotypes are present in poultry and poultry products. While serotypes such as *S. Enteritidis* and *S. Typhimurium* are extensively studied, many other poultry-associated serotypes remain largely unexplored in terms of their ecology and epidemiology. This knowledge gap is particularly concerning given the increasing prevalence of antimicrobial resistance among *Salmonella* strains isolated from US poultry. Addressing this issue requires continued surveillance and research to better understand the distribution and resistance patterns of lesser-known *Salmonella* serotypes [9-11].

To combat such pathogens, the poultry processing industry relies on various sanitation measures, including inorganic compounds based on chlorine-based products, and organic compounds based on quaternary ammonium compounds. Peracetic acid (PAA), a widely used organic sanitizer in industries such as healthcare, wastewater treatment, fine chemicals, and paper production [12], has emerged as a key disinfectant in poultry processing particularly in chiller tanks, as well as to disinfect equipment [13-16]. PAA disinfection not only requires maintenance of lower residual concentrations but also generates fewer disinfection by products [17-23]. It has also recently been found to exhibit greater antimicrobial efficacy as compared to chlorine dioxide and sodium hypochlorite against *Salmonella* Typhimurium on chicken skin and food-contact surfaces [24].

A crucial aspect in ensuring adequate residual disinfectant concentration to achieve bacterial inactivation while preventing excessive residual levels, involves evaluating how poultry chiller water characteristics influence PAA decay. However, studies in this area remain limited, with existing reports restricted to preliminary assessments. Previous studies have predominantly relied on microparameters such as chemical oxygen demand (COD) or dissolved organic carbon (DOC) to evaluate PAA degradation. Variations in organic matter levels influence PAA decay in water. However, COD is an unreliable parameter for assessing PAA decay in poultry processing water due to its inability to distinguish between reactive and non-reactive organic matter, its susceptibility to interference from inorganic substances, and its inconsistent correlation with PAA degradation rates [13]. Furthermore, COD measurement is labour-intensive and lacks the precision required for real-time monitoring. Therefore, our hypothesis was that total dissolved solids (TDS) would be a better indicator than COD for assessing PAA decay in poultry processing water. Unlike COD, TDS provides a more comprehensive and stable representation of PAA decay by accounting for both organic and inorganic dissolved substances, ensuring more reliable and repeatable measurements [25]. Its ability to offer real-time data makes TDS a superior metric for optimizing PAA dosing, enhancing disinfection efficiency and improving microbial safety in poultry processing environments.

There is a paucity of information on the fate of PAA in poultry processing chiller water systems and its possible effect on pathogen survival. Therefore, the objectives of this work were to (i) perform lab-scale pathogen shedding, transfer, and disinfection experiments during the chilling stage of poultry processing, with the goal of identifying optimal sanitization strategies; (ii) use data from these experiments to build mathematical models that can accurately predict results from the lab scale all the way to pilot and commercial scales, and iii) assess the efficacy and decay kinetics of PAA in simulated poultry processing environments, highlighting its interactions with organic matter and its potential to improve pathogen control.

## 2. Materials and Methods

### 2.1. Reagents

Technical grade peracetic acid (PAA) solution was supplied by Lab Alley (Spice Wood, TX, USA) and contained 15% of peracetic acid (PAA), 22% of hydrogen peroxide, 16% of acetic acid and the rest water. Separately, 25% hydrogen peroxide (H_2_O_2_) solution (Lab Alley, Spice Wood, TX), 99 % acetic acid solution (Sigma-Aldrich Inc., St. Louis, MO), chemical oxygen demand (COD) reactor (Lovibond RD125, MD100, Tintometer Inc, Sarasota, FL), Lovibond COD vials (Tintometer Inc.), Palintest Kemio Disinfection kit (Global Treat, Inc., Spring, TX), PAA high-range electrochemical sensors (KEM25PAA) and low-range sensors (KEM25PAL) (Global Treat, Inc., Spring, TX), a portable meter (Oakton, PCTSTestr^TM^ 50; Cole-Parmer, Vernon Hills, IL) to measure pH, TDS and conductivity, sodium hydroxide (Sigma-Aldrich Inc., St. Louis, MO), sodium bicarbonate (Sigma-Aldrich Inc.), lecithin (Thermo Fisher Scientific, Ward Hill, MA), buffered peptone water (BPW; Neogen, Lansing, MI), and a hydrogen peroxide (H_2_O_2_) test kit (Taylor K-1826, ChemWorld, Kennesaw, GA) were purchased from respective vendors.

### 2.2. Types of chicken samples tested

Two types of chicken samples were tested: non-frozen whole chicken carcasses and chicken thigh fryer pieces. Both chicken samples were purchased from a local grocery store, kept at 4 °C, and used for experiments on the day of purchase. Separately, non-frozen whole chicken carcasses (each weighing around 4 − 5 lbs) were purchased directly from a commercial poultry processing facility in Ohio, right before the carcasses entered the immersion chiller tank. For bacterial studies, chicken fryer thighs weighing 100 ± 20 g were used [13].

### 2.3. PAA decay and water chemistry in the presence of whole chicken carcasses

These experiments were conducted to study the decay of PAA in the presence of an organic load. The poultry chilling process was simulated using a bench-top immersion chiller tank containing 10 L of chilled (4°C), sterile, deionized (DI) water mimicking chiller water temperatures in commercial facilities. Two initial PAA concentrations were tested: 200 mg·L^-1^ and 70 mg·L^-1^. The pH was adjusted to around 7.2 in all experiments using 10 N sodium hydroxide. Two types of chicken carcasses were tested: local grocery store bought, and carcasses obtained from a commercial poultry processing facility. A whole chicken carcass (∼4 lb; with skin) was placed in the PAA-added chilled water tank, and required amount of water aliquots were taken every minute for 60 min, under manual stirring. The COD, TDS, conductivity, and pH levels were measured at 5-minute intervals. All experiments were conducted in triplicate. The plant-processed (pre-chill) carcasses were studied under similar experimental conditions. COD was determined using the thermoreactor digestion method [26], employing high-range vials at 150°C for 120 min. PAA levels were measured immediately after sampling using a high-range electrochemical sensor-based method with a Palintest Kemio Disinfection kit. The pH, TDS, and conductivity were continuously monitored using a digital Oakton portable meter, and DI water was used as control for all measurements.

### 2.4. Water chemistry in the presence of whole chicken but with no PAA

Local grocery store bought whole chicken carcasses (∼4 lbs) were tested in this study. The carcasses were immersed at 4°C in bench-scale 10 L immersion chilling tank. Aliquot samples from the chiller water were taken at regular intervals, under manual agitation to measure COD levels. At the same time points, TDS, conductivity, and pH were also measured. The total duration of each experiment was 60 min. All experiments were conducted in triplicate.

### 2.5. Acetic acid and hydrogen peroxide decay in the presence of whole chicken

Disinfectant decay experiments were performed in 10 L chilling tanks using 200 mg·L^-1^ PAA, as described earlier. When 200 mg·L^-1^ of PAA was added to a 10 L chiller tank, it decomposed into 213 mg·L^-1^ of acetic acid (AA) and 300 mg·L^-1^ of H_2_O_2_. Two separate experiments were done to study the individual decay dynamics of AA and H_2_O_2_ in the presence of an organic load. A total of 13.3 mL of concentrated (16%) AA and 13.64 mL of (22%) H_2_O_2_ were added separately to chilled water tanks to achieve initial concentrations of approximately 213 mg·L^-1^ AA and 300 mg·L^-1^ H_2_O_2_, respectively. The experimental conditions of the chiller water tank system, and the procedure employed, are described in section 2.3. After adding the store-bought whole chicken carcass (under manual agitation), aliquot samples from the chiller water were taken every 5 min and H_2_O_2_, conductivity, COD, TDS, and pH levels were measured over 1 h. H_2_O_2_ levels were measured immediately after sampling using vendor’s recommended protocols with a low-range TK3345-Z test kit (ChemWorld, Roswell, GA, USA). AA levels were assessed indirectly using real-time data from an immersed conductivity probe. All experiments were conducted in triplicate.

### 2.6. Salmonella cocktail preparation

Five serovars of *Salmonella enterica (S. Infantis, S. Enteritidis, S. Typhimurium, S. Newport, S. Javiana* were provided by Dr. Manan Sharma, Environmental Microbial and Food Safety Laboratory (USDA ARS NEA, Beltsville, MD). After opening, each overnight-grown strain was supplemented with 15% sterile glycerol and stored at −80 °C for long-term use. Prior to each experiment, a loop of frozen cultures from each strain was streaked onto blood agar plates (Tryptic Soy Agar (TSA) with 5% Sheep Blood; Thermo Fisher Scientific, Lenexa, KS) and incubated overnight at 37 °C. From these agar plates, a single colony from each strain was selected and cultured in individual 9 mL aliquots of Brain Heart Infusion (BHI) broth (Neogen, Lansing, MI) at 37°C overnight. From each individual 9 mL culture, a 1-mL aliquot of a *S. enterica* strain was pipetted and added to a 15 mL centrifuge tube, resulting in a total of 5 mL of the strain cocktail. This cocktail was enumerated using the serial dilution colony-forming unit (CFU/mL) plate count method on the day of each experiment. Serial dilutions were prepared in 1× nBPW, and colonies were enumerated by aliquot plating on brilliant green agar with sulfapyridine agar plates (BGS; Neogen). The BGS plates were then incubated overnight at 37°C to determine the initial bacterial load. To achieve a high (7 log CFU·10 μL^-1^) or low (5 log CFU·10 μL^-1^) bacterial load, the inoculum cocktail was further diluted with sterile 1× nBPW. This diluted cocktail was used on the same day to inoculate fresh store-bought chicken thigh pieces (100 ± 20 g) by applying 10 µL of cell suspension. The cell suspension was spread with a sterile glass rod over approximately 100 cm^2^ of the surface of the thigh pieces and allowed to attach for 10 min before starting the rinsing process.

### 2.7. PAA efficacy on S. enterica cocktail in the absence of organic load

This study focused on the mechanisms by which PAA inactivates a 5-strain *S. enterica* cocktail in the absence of an organic load. PAA was added to 100 mL of sterile DI water in a glass beaker to achieve 1, 3, 5, or 10 mg·L^-1^ residual PAA stock solutions. An appropriate volume of the cocktail was added to the beakers to yield high or low bacterial concentrations, respectively. Flasks were continuously gently stirred using an overhead stirring apparatus equipped with sterile paddles. At preset times (0.5, 1, 3, 5, and 10 min), 500 µL aliquots were sampled from the beakers and neutralized with 500 µL of nBPW. Bacteria survival was measured by counting colonies (after overnight incubation at 37 °C for 48 h) on BGS agar plates. All experiments were independently repeated three times.

### 2.8. Bacterial shedding from chicken thighs in the absence of PAA

Results of preliminary studies showed that a 10 min attachment period was the most optimal microbial load attachment time on chicken fryer thighs. Store-bought chicken thighs with skin (100 ± 20 g) were exposed to UV light for 10 min to eliminate any residual bacteria. These thighs were exposed to high or low levels of *S. enterica* cocktail inoculum for 10 min, after being spread using a sterile glass rod. Each chicken thigh piece was aseptically transferred into individual, sterile, 1.6 L homogenizer blender filter bags (Nasco, Whirl-Pak™, Pleasant Prairie, WI) containing 100 mL of sterile nBPW. The Whirl-Pak bags were shaken vigorously by hand for 1, 3, or 5 min, and the pieces were then aseptically transferred into a second nBPW rinse bag and shaken for another 1, 3, or 5 min. This process was repeated for a total of five rinses. In each step, a 10 µL aliquot of the rinse sample was diluted with sterile nBPW as needed, and then an aliquot was spread onto a BGS agar plate, which was incubated at 37 °C for 24 h for colony enumeration. CFUs were enumerated manually in each plate and CFU per thigh of the total rinse volume was calculated. Three replications were performed for each condition.

### 2.9. PAA efficacy on S. enterica cocktail in the presence of organic load

Chicken thigh samples were prepared and inoculated with the bacteria cocktail as described in section 2.8. For the first rinse of the thigh piece, the Whirl-Pak filter bags contained 100 mL of deionized water with initial PAA concentrations of 10, 20, 30, 40, or 50 mg·L^-1^. For the second rinse, the filter bags contained 100 mL of sterile nBPW with no PAA. The first rinse was done using a 1-, 2- or 3-min exposure time, while the second rinse in nBPW solution was done for 1 min. Aliquots from first rinse bags were neutralized with nBPW as needed and bacteria enumerated using methods previously described in section 2.7. All procedures were conducted in triplicate.

### 2.10. Mathematical model for PAA decay and pathogen inactivation

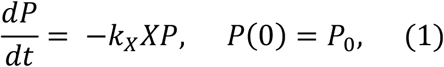

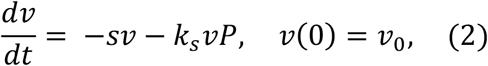

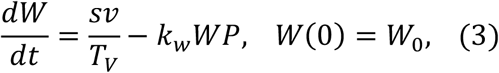

The decay rate of PAA (*P*(mg·L^-1^)) in chiller water at any time *T* is given by Equation (1), where *X* (mg·L^-1^) is the concentration of organic and inorganic matter released from chicken into the chilled tank, and *k*_*X*_(L·mg^-1^·min^-1^) is the apparent reaction rate constant for the reaction between PAA and *X* in the chiller water. Here, *X* can be represented by a measurable parameter such as COD or TDS, and will most likely be described by organic matter [27]. Equation (2) quantifies the change in bacterial levels, *v* (CFU) on the surface of chicken in the chiller tank, where *s* (min^-1^) represents the bacteria shed rate from the chicken surface into the chiller water and the second term describes the bacteria killing rate on the chicken surface via PAA, governed by *k*_*s*_ (L·mg^-1^ ·min^-1^). Equation (3) captures the change in bacterial levels, *W* (CFU·mL^-1^) in the chiller water in terms of (i) bacteria shed from the chicken surface to the water (*T*_*V*_is the water volume (mL)), and (ii) the pathogen killing rate in water via PAA, described by the term involving *kw*(L·mg^-1^ ·min^-1^).

#### 2.10.1. Mathematical form for s (shedding)

Results from a consecutive whole carcass rinse study by Lillard [2] tracking the recovery of aerobic bacteria and Enterobacteriaceae from chicken carcass surfaces, indicated that the total amount of bacteria shed after *n* consecutive (same duration) rinses are well described by a saturation function. In the equation below, *w* (CFU) represents the total amount of bacteria shed from a carcass after *n* consecutive rinses:

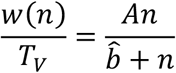

*A* (CFU·mL^-1^) approximates the total bacteria on the carcass that can transfer to the water and 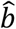 represents the number of consecutive washes needed for 50% of *A* to shed.

As we wanted *w* to be a continuous function of rinse time *T* (min), the following equation 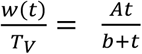 was used. It should be noted that 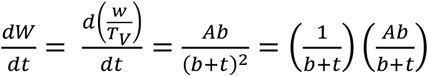, resembles the first term on the RHS of equation (3), where 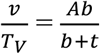 (the amount of bacteria shedding from the carcass at rinse time *T*). Thus, we quantified the shed rate 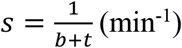, where *b* is the time necessary for 50% shedding to occur.

#### 2.10.2. Parameter determination and statistical analysis

Parameter estimation involving non-linear mathematical forms were done using the Levenberg-Marquardt (LM) algorithm [28]. Linear regression, ANOVA, and two-sample test were performed with standard packages in MATLAB version R2024a. Model (Eq 1-3) predictions were realized via the ode45 package in MATLAB. All experiments were repeated three independent times unless indicated otherwise and data reported as mean ± standard deviation.

## 3. Results and Discussion

### 3.1. PAA decay and water chemistry in the presence of fresh whole chicken carcasses

#### 3.1.1. Experimental results

Results from the whole chicken carcass exposed to 200 mg·L^-1^ or 70 mg·L^-1^ PAA in 10-L chilling tanks indicated that the organic matter from the carcasses released slowly into the tank, with consistent increase in COD and TDS levels over the 60 min period, for both plant bought and store bought chicken (**Fig. 1**). For all studies, COD and TDS levels in chilled DI water were measured before introducing PAA. At the start of the experiment (at 0 min), in tanks containing 200 mg·L^-1^ PAA, the initial COD level was 700 ± 20 mg·L^-1^ and TDS level was 250 mg·L^-1^ (**Fig. 1A**). Right after introducing the carcass (at 1 min) the COD slightly increased to 785 mg·L^-1^ for store bought chicken, and to 750 mg·L^-1^ for plant bought chicken. After 60 min, the final COD concentration for store-bought chicken was 1787.7 mg·L^-1^, and for plant bought chicken was 1560.5 mg·L^-1^. Similar increases in TDS levels were also noted after introducing the chicken. After 60 mins, TDS levels rose to 487.3 mg·L^-1^ for store-bought chicken, and to 422 mg·L^-1^ for plant-bought chicken.

**Fig. 1.**
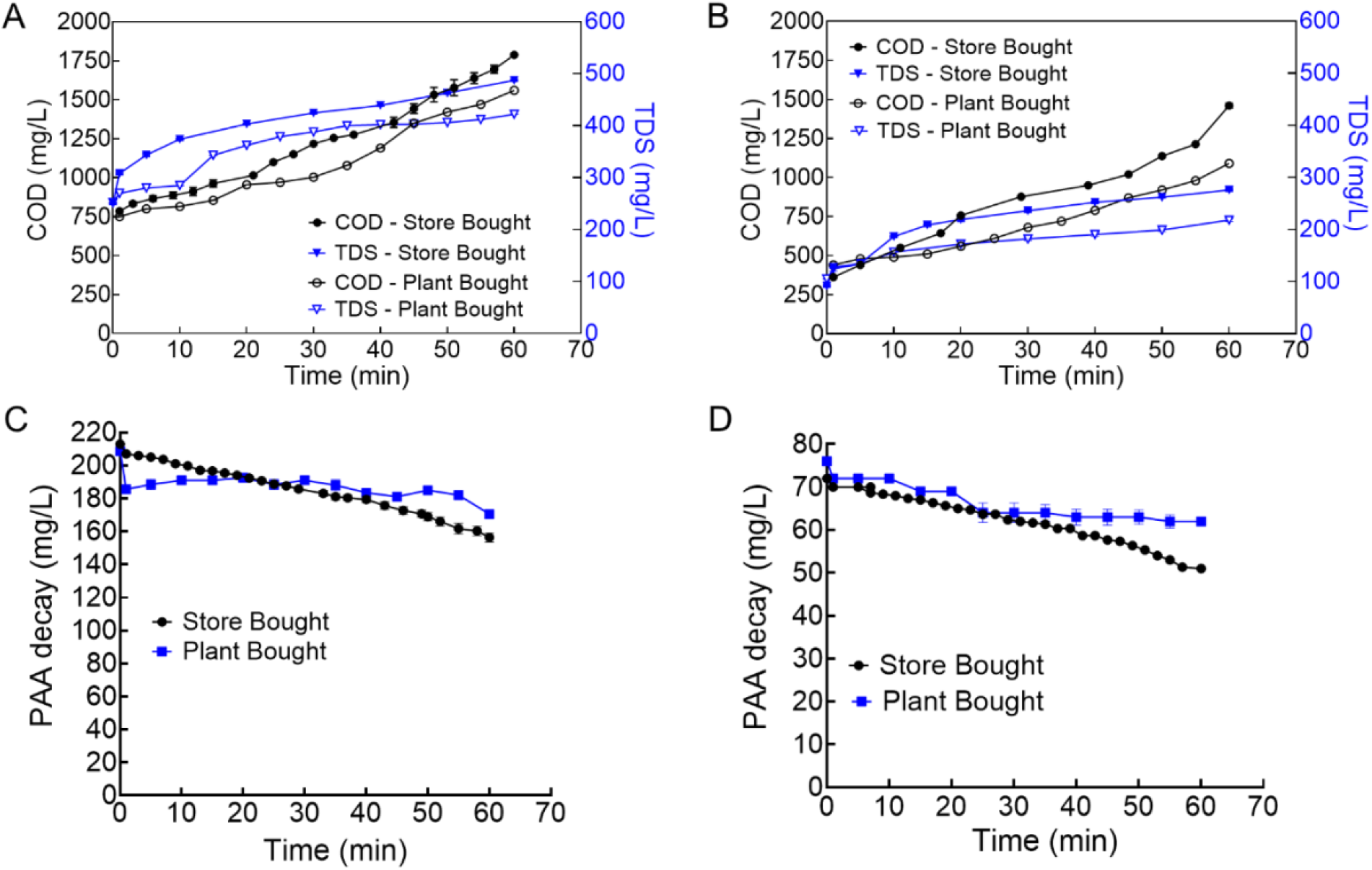
Water chemistry and PAA decay profiles in 10-L chilling tanks containing whole chicken carcass with skin. COD (black) and TDS (blue) levels in the tank containing store vs. plant bought chicken, exposed to initial PAA dose of 200 mg/L (**A**) or 70 mg/L (**B**). PAA decay in the chiller tanks containing store vs. plant bought chicken, from starting concentration of 200 mg/L (**C**) or 70 mg/L (**D**). Data presented were the mean ± standard deviation of 3 independent replicates for each condition.

In tanks containing 70 mg/L PAA, at the start of the experiment (at 0 min), the COD level was 304 mg·L^-1^ and TDS level was around 100 mg·L^-1^ (**Fig. 1A**). After introducing the carcass (at 1 min) the COD slightly increased to 361 mg·L^-1^ for store bought chicken, and to 440 mg·L^-1^ for plant bought chicken. After 60 min, the COD concentration for store-bought chicken was 1460 mg·L^-1^, and for plant bought chicken was 1090 mg·L^-1^. Similar increases in TDS levels were also noted after introducing the chicken. After 60 mins, TDS levels rose to 276 mg·L^-1^ for store-bought chicken, and to 218 mg·L^-1^ for plant-bought chicken. COD measures the amount of organic material that can be oxidized, while TDS quantifies the total concentration of dissolved solids in the water. The significant spike in COD and TDS levels at 0 min after the addition of PAA was due to the release of byproducts from PAA, i.e., acetic acid and H_2_O_2_. The relationship between COD and TDS demonstrated that the increase in dissolved solids was directly linked to the breakdown of organic material during the chilling process. The higher COD and TDS levels observed at 200 mg·L^-1^ PAA suggest that this concentration promoted more extensive organic matter breakdown, resulting in a greater release of dissolved solids. The larger increases in COD and TDS for the store samples suggest that they contained more organic material, which was broken down during the chilling process, contributing to greater levels of dissolved solids and oxidizable compounds in the chiller water.

We also measured the PAA decay in these chiller tanks, that were maintained at 200 mg·L^-1^ or 70 mg·L^-1^ concentration at the start of experiment. PAA decay was faster for store-bought chicken, at both PAA levels tested. After 60 min, the PAA concentration dropped to 156.3 mg·L^-1^ in the store-bought chicken tank, and to 166.5 mg·L^-1^ in the plant-bought chicken tank, when the initial concentration was 200 mg·L^-1^ (**Fig. 1C**). Similarly, PAA concentration dropped to 51 mg·L^-1^ in the store-bought chicken tank, and to 62 mg·L^-1^ in the plant-bought chicken tank, when the initial concentration was 70 mg·L^-1^ (**Fig. 1C**), at the end of 60 min. The faster PAA decay for store-bought samples can be attributed to the higher organic load, which increases the demand for PAA to oxidize organic material.

The exudates and debris from poultry carcass in PAA chiller water also affect the pH. In general, the pH of chilled DI water containing PAA ranged between 3 to 4, before adjustment. The pH was relatively stable during immersion chilling, with a slight decrease from 7.5 (after adjusting with NaOH) to 6.5 for all conditions tested. This pH was in the range recommended for maximizing the efficacy of PAA concentration against microorganisms. No significant changes in our model parameters were noted due to this pH change.

#### 3.1.2. Modeling results for PAA decay dynamics

##### 3.1.2.1. PAA decay (P_0_ = 200, 70) in 10-L tanks with store & plant bought whole carcasses

To predict PAA levels in the water during chilling, it is necessary to determine the rate at which PAA decays in the presence of carcass exudates and constituents at 4× C. The objective was to test which proxy indicator – COD or TDS – would be a better predictor of PAA levels under simulated chiller conditions. Utilizing equation (1) and TDS/COD and PAA levels from whole carcass batch experiments, and the LM algorithm [MATLAB], we fit the apparent reaction rate constant for the decay of PAA and in the chiller water given by *k*_*X*_(L·mg^-1^·min^-1^), where *X* represented either TDS or COD. Results were presented in **Table 1** for store-bought and plant-bought (pre-chiller carcasses from a commercial plant) chicken. At initial PAA concentrations of 70 ppm and 200 ppm, *k*_*TDS*_provided more consistent and accurate fits with respect to store and plant chicken when compared with *k*_*COD*_(Table 1). These findings suggest that TDS information predicts PAA decay better than COD in the context of poultry chilling.

**Table 1.** Results from the model fit to experimental data obtained from store-bought and plant-bought chicken.

| | PAA<br>($\text{mg}\cdot\text{L}^{-1}$) | TDS | | COD | |
| --- | --- | --- | --- | --- | --- |
| | | $k_{TDS}$ | RMSE<br>($\text{mg}\cdot\text{L}^{-1}$) | $k_{COD}$ | RMSE<br>($\text{mg}\cdot\text{L}^{-1}$) |
| Store-bought | 70 | $3.6 \pm 0.3\text{E-}5$ | $1.2 \pm 0.2$ | $1.2 \pm 0.04\text{E-}5$ | $1.8 \pm 0.1$ |
| | 200 | $1.7 \pm 0.6\text{E-}5$ | $3.4 \pm 1.2$ | $1.29 \pm 0.4\text{E-}5$ | $8.3 \pm 3.2$ |
| Plant-bought | 70 | $3.9 \pm 2.1\text{E-}5$ | $3.2 \pm 2.3$ | $1.7 \pm 0.9\text{E-}5$ | $5.1 \pm 4.0$ |
| | 200 | $1.5 \pm 0.7\text{E-}5$ | $7.6 \pm 7.4$ | $0.9 \pm 0.3\text{E-}5$ | $9.2 \pm 8.9$ |

##### 3.1.2.2. *Using TDS vs COD to predict PAA fate in commercial pre-chiller & chiller operati*ons

To test the hypothesis that TDS information is a better predictor of PAA levels than corresponding COD data, we adapted equation (1) to apply to a commercial chiller process with continuous PAA input as well as water inflow and outflow as follows:

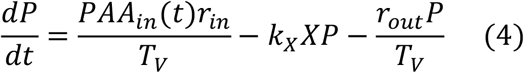

In equation (4), *P*(mg·L^-1^) is the concentration of PAA in the chiller water, *PAA*_*in*_(*T*) (mg·L^-1^) is the input PAA concentration, *r*_*in*_(L·min^-1^) is the average flow rate of the water/PAA mixture entering the chiller, *r*_*out*_(L·min^-1^) is the average flow rate of water leaving chiller, *T*_*V*_ (L) is the tank volume, *X* (mg·L^-1^) is the TDS or COD level, and *k*_*X*_ represents the apparent 2^nd^ order rate constant of PAA with material in the chiller water (*k*_*X*_ = *k*_*TDS*_ or *k*_*COD*_, when *X* = TDS or COD respectively).

Equation (4) was applied to an industrial setup (pre/main chiller processes) in a typical high-speed poultry processing plant in North America. The tank volume for the pre-chiller was *T*_*V*_ = 27,000 (L) and for the main chiller was *T*_*V*_ = 35,000 (L). The carcass line speed was about 60 chickens per minute. Sampling of pre/main chiller water occurred over three days during routine operation with PAA used as disinfectant [29]. Chiller samples were aseptically collected by removing 125 mL of water from different areas of the chiller tank at pre-determined time points. PAA input concentrations as well as TDS, COD (2 days only), pH, temperature and PAA levels in both pre and main chiller processes were determined. Main chiller pH values were 7.0 ± 0.2, 7.0 ± 0.2, and 6.8 ± 0.1 over the respective 3 processing days. The main chiller water temperatures were 1.9 ± 0.5 °C, 2.3 ± 0.5 °C, and 2.6 ± 0.2 °C, respectively; pre-chiller pH values were 7.2 ± 0.2, 7.2 ± 0.1, and 7.2 ± 0.09, respectively; and pre-chiller water temperatures were 2.7 ± 0.4 °C, 3 ± 0.5 °C, 3.0 ± 0.6 °C, respectively. For processing days 1 and 2, the incoming PAA concentrations for the pre-chiller varied between 27≤ *PAA*_*in*_(*T*) ≤ 35 (mg·L^-1^) and 26 ≤ *PAA*_*in*_(*T*) ≤ 32 (mg·L^-1^), respectively. For processing days 1 and 2, the incoming PAA concentrations for the main chiller varied between 60 ≤ *PAA*_*in*_(*T*) ≤ 65 (mg·L^-1^) and 49 ≤ *PAA*_*in*_(*T*) ≤ 65 (mg·L^-1^), respectively. For both processing days, the flow rates in and out (*r*_*in*_ and *r*_*out*_) varied between 75 to 90 (L·min^-1^) for the pre-chiller and between 95 to 115 (L·min^-1^) for the main chiller, respectively. Using data from processing day 1, model equation (4) and the LM algorithm [MATLAB], we determined the optimal fits for *k*_*TDS*_ and *k*_*COD*_ in both the pre-chiller and main chiller contexts. To illustrate the utility of the model, we used the respective fit values for *k*_*TDS*_ and *k*_*COD*_ to predict the PAA dynamics in the pre-chiller and main chiller tank during processing day 2. **Figure 2** below shows the prediction results against the commercial data on processing day 2.

**Fig 2.**
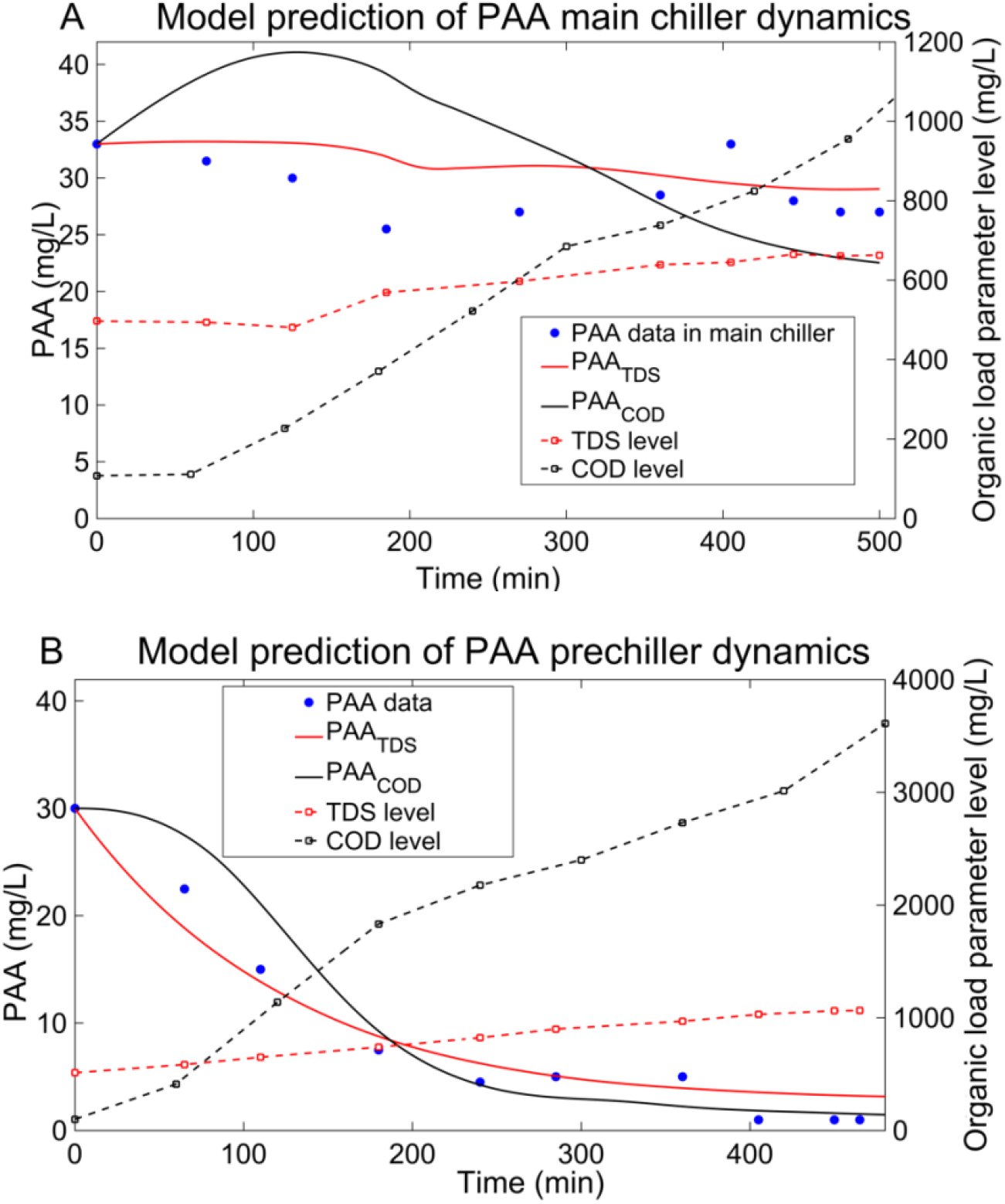
Model prediction of PAA dynamics in a commercial pre-chiller/main chiller setup during processing day 2 [29]. Model fit to PAA and TDS/COD data from processing day 1 in both tanks, obtaining *k*_*TDS*_ = 1.45E-5 (L·mg^-1·^min^-1^) and *k*_*COD*_ = 1.04E-5 (L·mg^-1·^min^-1^) for the pre-chiller and *k*_*TDS*_ = 6.1E-6 (L·mg^-1·^min^-1^) and *k*_*COD*_ = 1E-5 (L·mg^-1·^min^-1^) for the main chiller, respectively. **A**. Processing Day 2 pre-chiller PAA prediction using TDS and COD levels. RMSE = 1.89 mg·L^-1^ for TDS and RMSE = 2.51 mg·L^-1^ for COD. **B**. Processing Day 2 main chiller PAA prediction using TDS and COD levels. RMSE = 3.09 mg·L^-1^ for TDS and RMSE = 6.06 mg·L^-1^ for COD.

PAA dynamics were also predicted during processing day 3 in terms of TDS measurements (note COD was not measured for this processing day). In the pre-chiller, the PAA model prediction yielded a root mean square error (RMSE) of 0.6695 (mg·L^-1^) and in the main chiller a RMSE of 5.433 (mg·L^-1^) over 8 h of processing. The flow rates were around 100 L·min^-1^ for first few hours of the shift and then increased to around 150 L·min^-1^ for the rest of shift. For the main and pre-chiller, PAA input concentrations varied from 35 mg·L^-1^ to 52 mg·L^-1^, and 28 mg·L^-1^ and 30 mg·L^-1^, respectively. Even with a 50% increase in water flow rates for the latter duration of the processing shift, the PAA dynamics model (equation 4) predicted PAA level changes accurately, as shown in Figure 2 A.

These prediction results indicate that TDS information more consistently and accurately predicted PAA dynamics rather than using COD data. However, using TDS to predict PAA dynamics is context specific (different *k*_*TDS*_ values across results as shown in Table 1 and Fig. 2). This indicates that model training is needed to determine *k*_*TDS*_ relative to specific chiller process specifications, which is relatively easy as TDS can be measured effectively in real time. Once the parameter *k*_*TDS*_ is determined, the model, along with calibration of the commercial system, can appropriate real time TDS as a compliance/verification check for PAA control during chilling.

Furthermore, rounding to the nearest order of magnitude, *k*_*TDS*_ and *k*_*COD*_ values determined from whole carcass in 10-L chiller tank experiments were similar to the *k*_*TDS*_ and *k*_*COD*_ values determined at the commercial scale (with continuous flow dynamics). That is, the time scale for PAA decay differed by less than an order of magnitude between the lab and commercial context. Considering the significant scaling differences between a lab simulated chiller process and a commercial process, this emphasizes the utility of lab experiments to capture fundamental dynamics of PAA fate during poultry chilling and further justifies experiments at the lab scale to determine inactivation rates of bacteria relative to PAA sanitation when PAA decay occurs.

### 3.2. Water chemistry in the presence of fresh whole chicken carcasses and no PAA

Store-bought whole chicken carcasses with skin were immersed in a 10-L chilled deionized water tank maintained at 4 °C for 60 min, without any PAA, to assess the baseline release of organic and dissolved solids. COD was measured to capture the complete oxidation demand over time. The initial COD levels began at 196 mg·L^-1^ at the 1 min timepoint, and reached a maximum of 897 mg·L^-1^ by 60 min. The increasing COD values, especially at later time points, indicated continuous leaching of oxidizable substances such as blood residues, lipids, and tissue debris from the carcasses, intensified by the presence of skin. Concomitantly, TDS levels increased from an initial 1.8 mg·L^-1^ at 0 min to 21.05 mg·L^-1^ at 1 min, and reached 144.5 mg·L^-1^ by 60 min. The sharp increase in the first 20–30 min suggests rapid early-phase leaching of soluble ions and low-molecular-weight compounds. The consistent rise in both COD and TDS throughout the 60-min period confirmed that whole chicken carcasses with skin contribute substantial organic and dissolved matter to chiller water, even in the absence of any chemical treatment. These findings establish a critical reference baseline for assessing the oxidative and solid burden introduced by untreated poultry during immersion chilling, highlighting the need for intervention strategies to manage water quality in commercial processing environments.

In the experiments without PAA, the pH and conductivity of the chilled deionized water were also monitored. The pH of the DI water (initially ∼7.0) dropped to a pH value of 6.1 over the 60 min chilling period, likely due to the release of organic acids and metabolic byproducts from carcass tissues and blood. Conductivity measurements similarly showed a substantial increase, increasing from 1.9 µS·cm^-1^ in DI water to 30.1 µS·cm^-1^ shortly after carcass addition, and a further steady increase to 213 µS·cm^-1^ by 60 min. This increase likely reflects the accumulation of ions and water-soluble compounds such as sodium, potassium, and chloride from carcass exudate proteins, as well as glycerides and surface debris. These shifts in pH and conductivity indicate that even in the absence of chemical disinfectants, the immersion of poultry carcasses significantly alters the chemical properties of chiller water, emphasizing the need for proper water quality management in processing systems.

### 3.3. Mechanisms of PAA decay in the presence of whole chicken carcasses

The addition of PAA to water results in its dissociation into acetic acid and hydrogen peroxide. To understand the underlying mechanisms by which such dissociated compounds contribute to the changes in TDS and COD, and which of them is predominant, we supplemented 300 mg·L^-1^ H_2_O_2_ or 213 mg·L^-1^ acetic acid (AA) to 10-L chiller tanks containing store-bought chicken. Results from these studies indicate that the addition of these compounds to the chiller water influenced both the release of organic matter and the oxidant decay patterns over time. A consistent increase in COD and TDS levels was observed over the 60 min (**Fig. 4**).

**Fig. 3.**
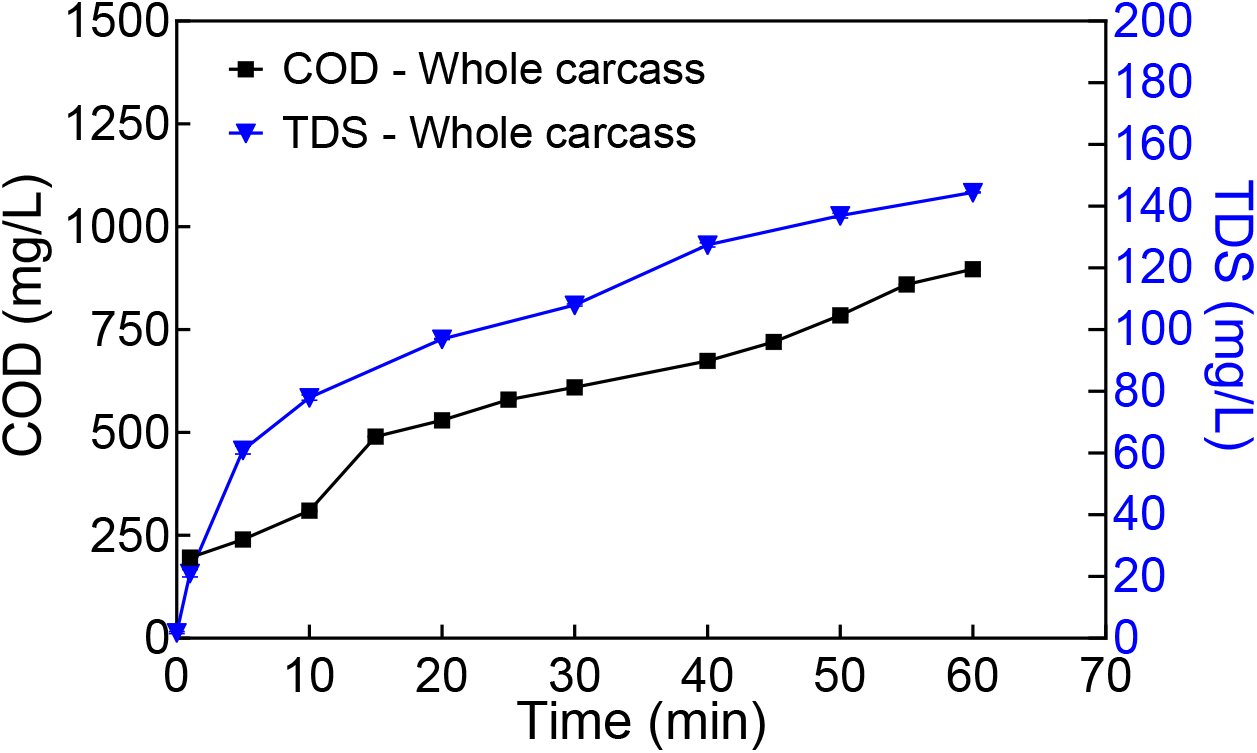
Water chemistry profiles in 10-L chilling tanks containing whole chicken carcass with skin. COD (black) and TDS (blue). Data presented were the mean ± standard deviation of 3 independent replicates for each condition.

**Fig. 4.**
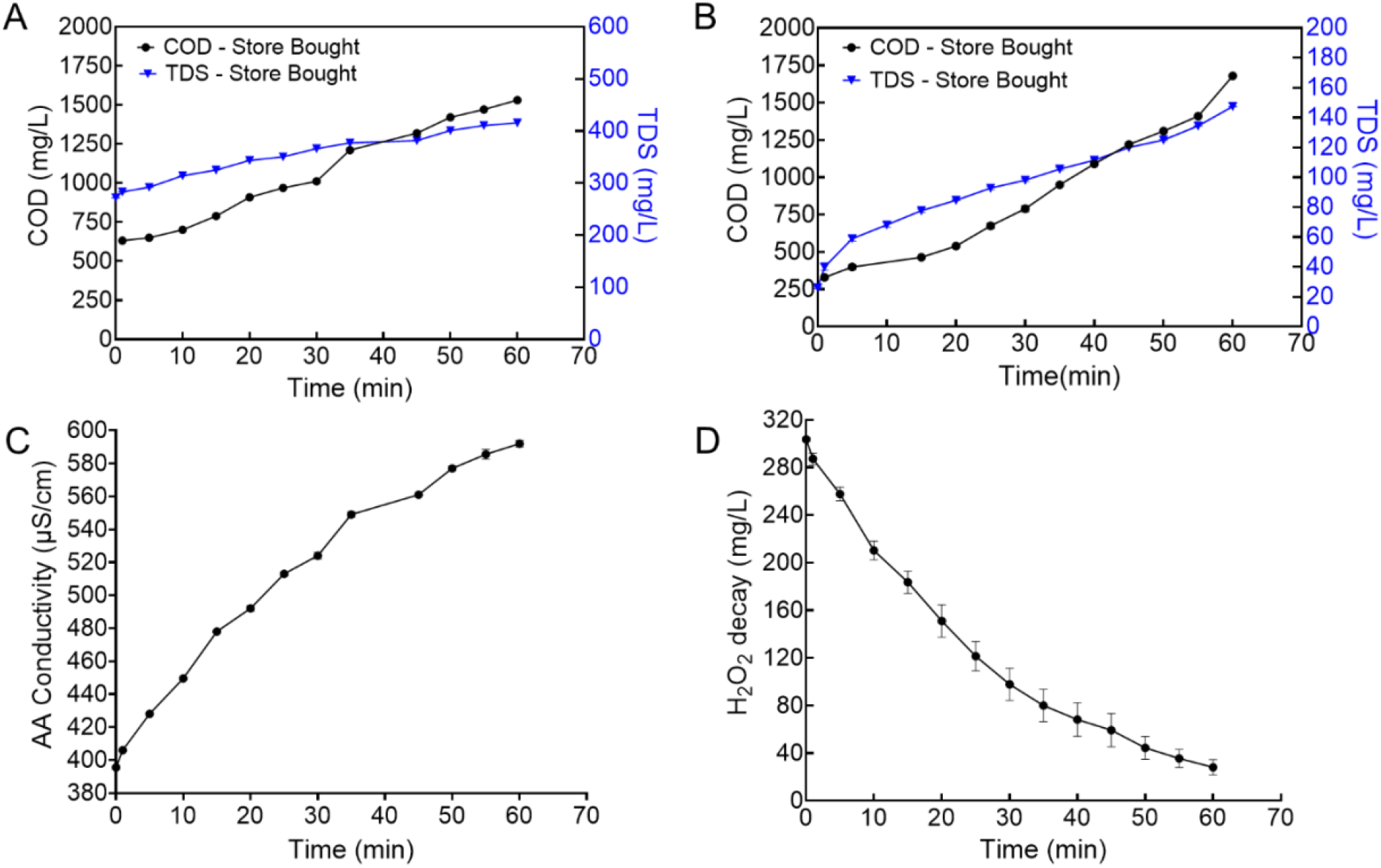
Water chemistry and H_2_O_2_ decay profiles in 10-L chilling tanks containing whole chicken carcass with skin. **A:** COD (black) and TDS (blue) levels in the chiller tank containing store-bought chicken, exposed to initial acetic acid (AA) dose of 213 mg·L^-1^. **B:** COD (black) and TDS (blue) levels in the chiller tank containing store-bought chicken, exposed to an initial H_2_O_2_ dose of 300 mg·L^-1^. **C:** AA conductivity increases in the chiller tank containing store bought chicken. **D**. H_2_O_2_ decay in the chiller tank containing store bought chicken. Data presented were the mean ± standard deviation of 3 independent replicates for each condition.

In the H_2_O_2_ treatment group, COD levels rose from 330 mg·L^-1^ at 0 min to 1680 mg·L^-1^ at 60 min, whereas TDS levels rose from 30 mg·L^-1^ at 0 min to 147.5 mg·L^-1^ by 60 min (**Fig. 4, A**). In the AA treatment group, COD levels rose from 520 mg·L^-1^ at 0 min to 1531 mg·L^-1^ at 60 min, whereas TDS rose from 272 mg·L^-1^ at 0 min to 415 mg·L^-1^ by 60 min (**Fig. 4, B**). Although the COD rose significantly faster in the H_2_O_2_ group at each time point, the TDS levels remained significantly lower than those observed in the AA group at each time point. Results indicate that (i) both AA and H_2_O_2_ supported organic matter oxidation as evident from rising COD and TDS levels, (ii) organic matter breakdown in the presence of AA resulted in a moderate but steady accumulation of both oxidizable and dissolved solids in the chiller tank, (iii) AA promotes the release of more soluble breakdown products, whereas H_2_O_2_ aggressively oxidizes particulate organic matter, and (iv) H_2_O_2_ decayed significantly faster than AA, suggesting greater oxidative reactivity but shorter stability in the presence of high organic loads.

In the above experiments, the pH and conductivity of the water were also monitored. With the introduction of organic content in the chiller tank, a slight decrease in pH was noted. Conductivity measurements were used as an indirect assessment of acetic acid levels in the chiller tank, and the rise in hydrogen ion concentration indirectly indicates the decomposition of AA (**Fig. 4, C**). At 0 min, the conductivity of pH-adjusted DI water containing 213 mg·L^-1^ AA was 390 µS·cm^-1^, which rose to 406 µS·cm^-1^ shortly after carcass addition, and continued to increase steadily to 590 µS·cm^-1^ by 60 min. Meanwhile, in the 300 mg·L^-1^ H_2_O_2_ supplemented chiller tank, the conductivity rose from 40 µS·cm^-1^ at 0 min to 214 µS·cm^-1^ by 60 min.

The H_2_O_2_ decay profile demonstrated rapid loss of oxidant activity over time (**Fig. 4, D**). Starting at 303.5 mg·L^-1^ at 0 min, the concentration dropped to 28.1 mg·L^-1^ by 60 min. At the end of the 70 min, only 10.4 mg·L^-1^ of residual H_2_O_2_ remained. The steep decay profile reflects the high reactivity of hydrogen peroxide with organic material released from the chicken carcasses during immersion, especially within the first 20 min of contact. Taken together, these findings highlight the contrasting dynamics of AA and H_2_O_2_ as oxidants and their respective impacts on chiller water chemistry during the immersion processing of poultry carcasses.

### 3.4. Salmonella shed from chicken thighs in the absence of PAA

#### 3.4.1. Experimental results

These studies were done on chicken thighs and not whole carcasses. Preliminary studies showed that in the absence of PAA, *Salmonella* cocktail shedding was not influenced by attachment duration, regardless of the bacterial concentration or the absence of skin (**Supplementary Figure 1**). Therefore, a 10-min bacterial attachment was chosen for all studies moving forward. For shedding studies, two different initial loadings of *Salmonella*, i.e., around 3.4×10^7^ CFU per 10 µL (high load) and 3×10^5^ CFU per 10 µL (low load) were tested on chicken thighs (**Fig. 5**). After a 10-min attachment at room temperature, the thigh pieces were shaken in Whirl-Pak bags containing nBPW for multiple rinses, each rinse lasting 1 min, 3 min, or 5 min. As shown in **Fig. 5**, at both high and low loads, bacterial shedding from chicken thighs was predominantly influenced by both the rinse time and number of rinses. Most of the bacteria (> 60%) were removed during the first rinse, with significantly less bacterial shedding occurring in subsequent rinses. The extended shaking time also improved bacterial removal consistency across later rinses compared to the 1-min shaking time. While the first rinse consistently accounted for most of the bacterial shedding across all shaking times, the impact of extended shaking was most evident in the second rinse, where prolonged shaking facilitated the removal of more strongly adhered bacteria. These results indicate the effectiveness of shaking for at least a minute to dislodge adhered bacteria and thus maximize bacterial removal during poultry processing.

**Fig. 5.**
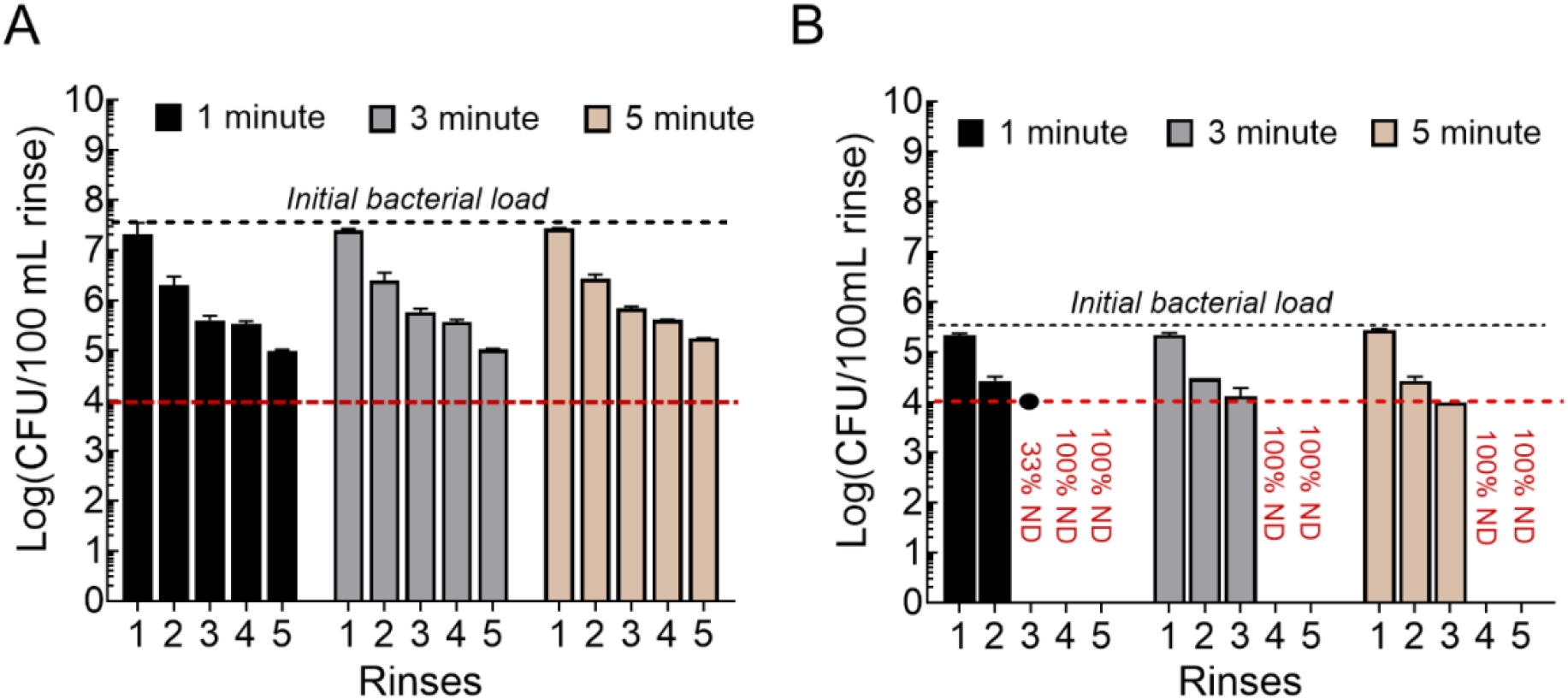
*Salmonella* shedding from chicken thighs in the absence of PAA, at (**A**) high initial bacterial load and (**B**) low initial bacterial load. The duration of the rinses and number of rinses were treated as variables to see their effect on bacterial detachment. The red dotted line represents the limit of detection. Solid circles in (**B**) illustrate the detected values while the percentage values given under the limit of detection line indicate the proportion of non-detected samples under those conditions.

#### 3.4.2. Modeling results for shedding

High and low inoculum shedding experiments were conducted using 5 consecutive wash times of 1 min each, 3 min each and 5 min each, respectively. As each inoculum level and wash time combination was run in triplicate, there were a total of 18 runs. The following equation was used:

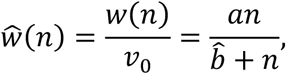

where *w* (CFU) was the total amount of bacteria shed from a carcass after *n* consecutive rinses and *v*_0_ (CFU) was the inoculum level on the chicken surface.

Using the LM algorithm [MATLAB] with data from the shedding experiments (see Appendix **section A**), we determined parameters *a* = 1.03 ± 0.06 and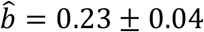, where errors represent the respective 95% confidence intervals. The motivation for considering rinse numbers only (and not rinse duration) comes from the results in Figure 5A and B which were not significantly different when comparing across consecutive rinse numbers. This means that 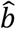 depends only on the consecutive rinse number as opposed to the rinse duration (in the context of 1, 3- and 5-min rinse times). This result also suggests that to determine the shed rate time scale, the shortest wash duration (1 min) should provide the most accurate approximation. Considering *w* as a continuous function of total rinse time, where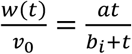, and fit for *b*_1_ with respect to the data from 1 min rinse duration, we found that *b*_1_ = 0.23 ± 0.11 (min). The fact that 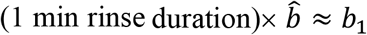 together with the fit results for 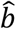 above, gave strong evidence that the time scale for 50% of the bacteria to shed from the chicken surface is 0.23 ± 0.04 min. Therefore, the *Salmonella* shed rate was set to 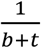, where *b* = 0.23 ± 0.04 min and *T* (min) is the total rinse time. This also indicates that shed rate does not depend on the inoculum load (high vs low) as both data sets (illustrated in Figure 5A and B) were used in determining *b*. Refer to Appendix **section A** for a typical fitting result for the model form 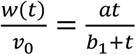 as well as an explanation as to how the non-detected samples from the low initial load rinse experiments (refer to Figure 5(B)) were incorporated in the parameter fit for *b*.

### 3.5. Salmonella survival in the presence of residual PAA and the absence of organic matter

#### 3.5.1. Experimental results

These experiments evaluated the bacterial inactivation efficiency of residual PAA solutions (1 – 10 mg·L^-1^) on *Salmonella* in the absence of an organic load. Two bacterial loads were chosen – high (2 × 10^7^ CFU·10 µL^-1^), and low (5 × 10^5^ CFU·10 µL^-1^) loads exposed to 30 sec, 1 min, 3 min, 5 min and 10 min. The results demonstrated a clear PAA dose and exposure time dependent reduction in bacterial counts. At a high load of *Salmonella*, bacteria did not survive at ≥ 3 mg·L^-1^ PAA after 3 min exposure, and ≥ 1 mg·L^-1^ PAA exposure beyond 10 min. Similar trends were noted at a low load of *Salmonella*, albeit even at lower exposure times, suggesting that higher PAA concentrations significantly enhance bacterial killing efficiency and reduced the required contact time for complete inactivation. These findings are particularly relevant for applications requiring sterilization in low-contaminant environments, where the absence of organic load ensures maximum residual PAA effectiveness. While lower concentrations (1 mg·L^-1^) may suffice given sufficient exposure time, higher concentrations (≥ 5 mg·L^-1^) are essential for rapid and complete bacterial inactivation within shorter timeframes. Optimizing residual PAA concentration and exposure time can, therefore, provide an effective strategy for bacterial control in chiller processing environments.

#### 3.5.2. Modeling Salmonella inactivation in chiller water (no organic load)

For these experiments, we used equation (3), with the shedding term removed (first term), as the killing rate of *Salmonella* in chilling water without any poultry/organic load was to be determined. So, equation (3) was adapted as follows:

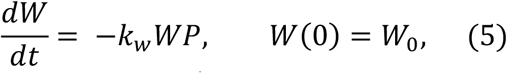

where *W*_0_ = initial inoculum level (CFU·mL^-1^). This is the Chick-Watson disinfection model [30] It should be noted that given the time scale of experiments (at most 5 – 10 min), the PAA concentration *P*= *P*_0_ (mg·L^-1^) remains as the initial PAA concentration set for each run. Equation (5), was then adapted as follows:

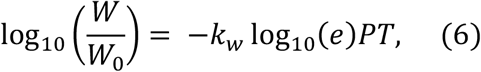

where 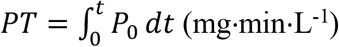.

Assuming that *T*^*^(min) is the time it takes for the bacterial concentration to become undetectable, equation (6) was modified to be:

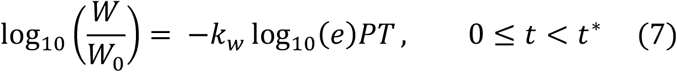

Using equation (7) and data from (**section 3.5.1**), the inactivation rate was calculated at various *W*_0_ and *P*_0_ and determined to be *k*_*w*_ = 0.15 ± 0.02 (L·mg^-1^·min^-1^) (**Supplementary Figure 2**). The goodness of fit was presented in Appendix **section B** using residual plot. ANOVA analysis suggested that there was not enough evidence to support the dependence of the inactivation fraction, 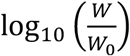, on the initial inoculum (*W*_0_) or initial PAA concentration (*P*_0_) (see Table in Appendix **section B**).

### 3.6. Killing rate of Salmonella in the presence of organic matter and PAA

#### 3.6.1. Experimental results

This study evaluated the bacterial inactivation efficiency of PAA solutions (10 – 75 mg·L^-1^) on *Salmonella* in the presence of an organic load. Two bacterial loads were chosen: high (1.2 × 10^7^ CFU/10 µL; **Fig. 7**), and low (5 × 10^5^ CFU/10 µL; **Fig. 8**) loads, exposed to 1 min, 2 min, or 3 min. The 10 – 75 mg·L^-1^ PAA values are initial concentrations, and they decayed very fast for all of the 1-, 2-, or 3-min experiments. Two rinses were considered: the PAA rinse represents the bacteria remaining in the rinse solution after 1 min PAA exposure, while the nBPW rinse reflects the bacteria that remained attached to the chicken thigh pieces after 1 min of PAA exposure and subsequent transfer to the neutralizing solution. For the high bacteria load, at 75 mg·L^-1^ initial PAA, bacteria survival was not observed and so the data was not shown in the plots. Results demonstrate a clear PAA initial dose and exposure time dependent reduction in the bacterial counts from both initial bacterial loads. In the PAA rinse step, bacterial survival decreased as the initial PAA concentration increased. Even after the nBPW rinse step, bacterial survival on the chicken thigh pieces was strongly influenced by the initial PAA concentration. Similar trends were noted for 2-min and 3-min exposure studies in the presence of high bacterial load. At the low bacterial load and 1-min exposure time (**Fig. 8**), complete bacterial inactivation (below detection level) was noted at an initial PAA level of ≥ 40 mg·L^-1^. However, at the 2-min exposure time, complete bacterial inactivation (below detection level) was observed with an initial PAA of ≥ 30 mg·L^-1^.

**Fig. 6.**
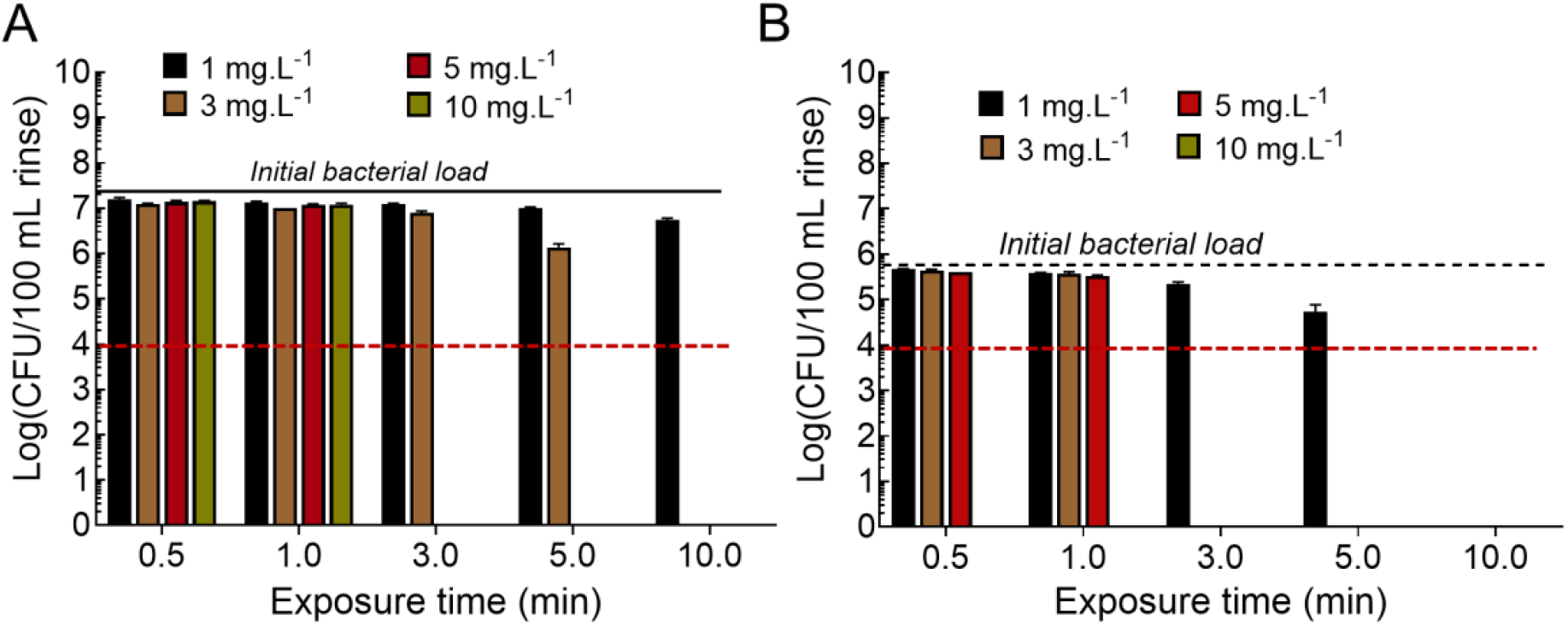
*Salmonella* survival in the presence of residual levels of PAA and the absence of organic matter, at (**A**) high initial load (2 × 10^7^ CFU·10 µL^-1^) and (**B**) low initial load (5 × 10^5^ CFU·10 µL^-1^). The red dotted line represents the limit of detection and conditions where all samples were below this limit were not represented by a bar here.

**Fig. 7.**
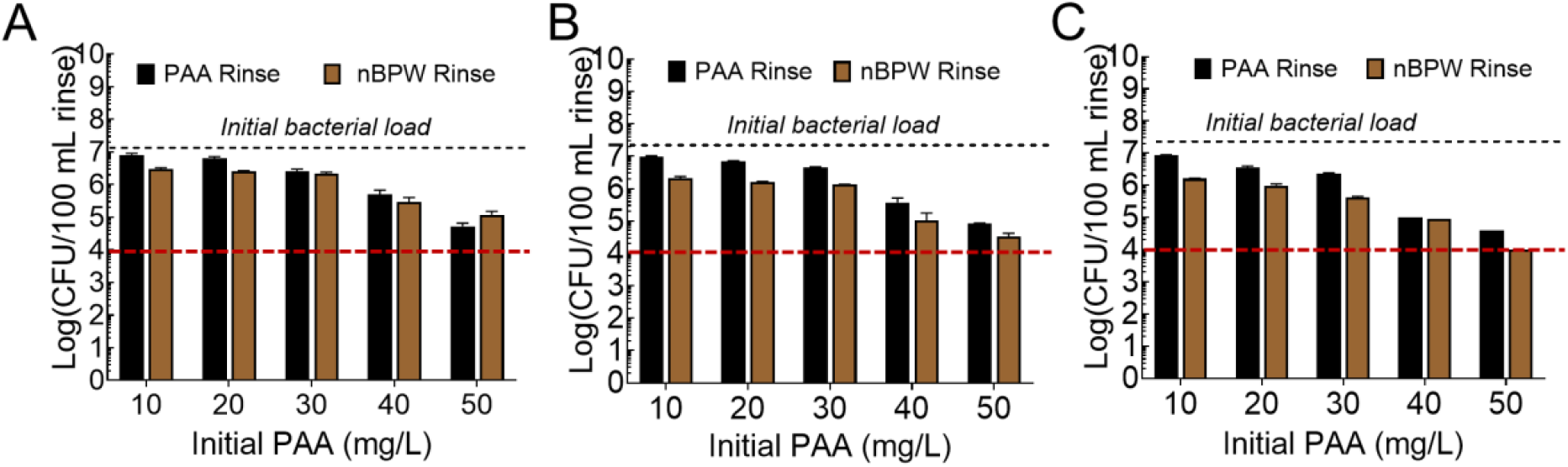
*Salmonella* survival in the presence of 10-50 mg·L^-1^ of initial PAA and organic matter, at a high initial bacterial load (∼1.2 × 10^7^ CFU·10 µL^-1^). After exposure to PAA (first rinse), the chicken thighs were rinsed with nBPW (second rinse), with each rinse lasting **(A)** 1, **(B)** 2, or **(C)** 3 min. PAA levels of ≥ 75 mg·L^-1^ completely inactivated the bacteria. The red dotted line represents the limit of detection.

**Fig. 8.**
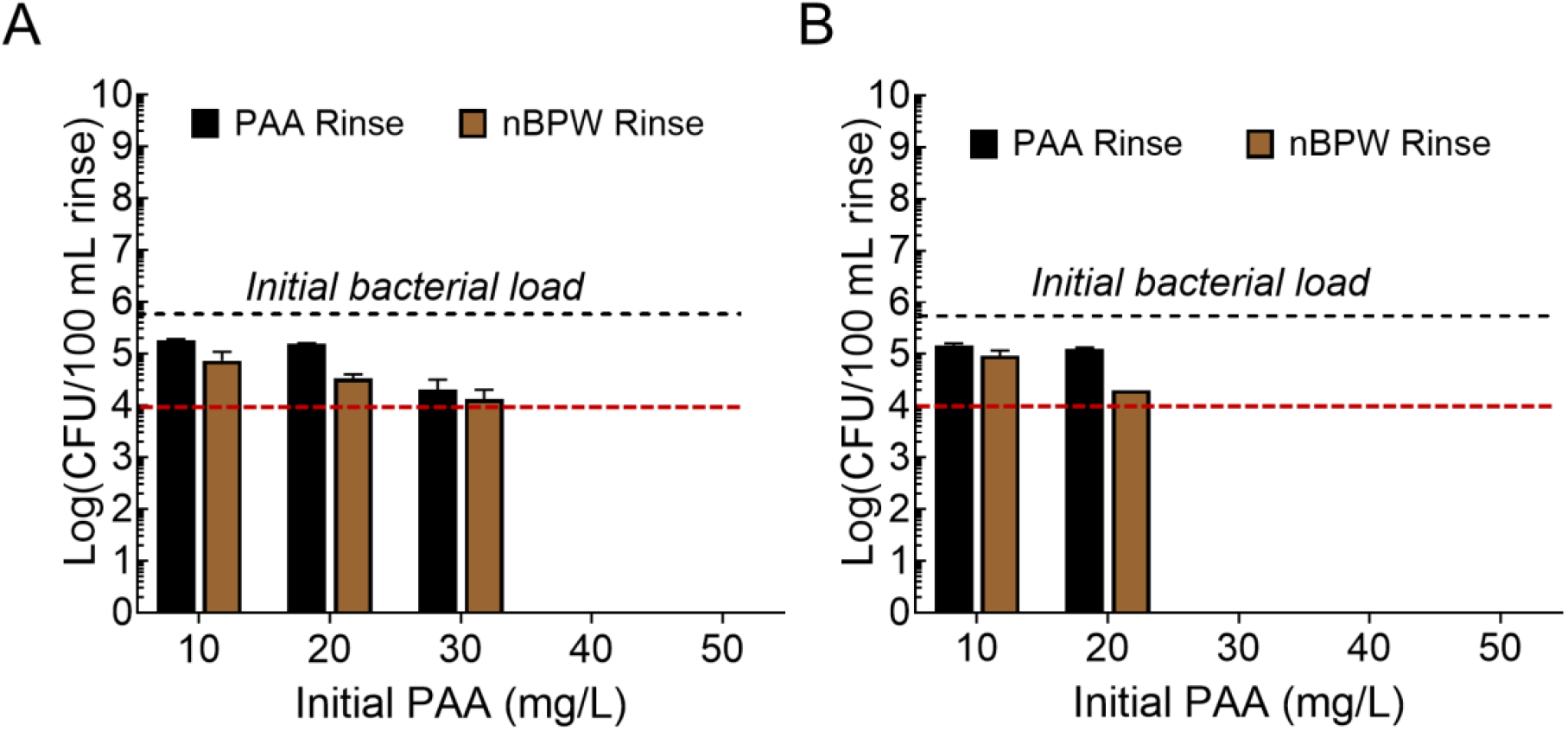
*Salmonella* survival in the presence of 10-50 ppm initial PAA and organic matter, at low load (∼5 × 10^5^ CFU·10 µL^-1^). After exposure to PAA (first rinse), the chicken thighs were rinsed with nBPW (second rinse), with each rinse lasting **(A)** 1, or **(B)** 2 min. The red dotted line represents the limit of detection.

These latter experiments demonstrated the critical role of both initial PAA concentration and exposure time in achieving effective *Salmonella* inactivation. Across 1-, 2-, and 3-min exposure times, bacterial reductions increased substantially at higher initial PAA concentrations. While lower initial concentrations (10–20 mg·L^-1^) provided moderate reductions, substantial bacterial survival was observed in both the PAA rinse and on chicken thighs. In contrast, initial PAA levels of ≥ 40 mg·L^-1^ consistently achieved > 99.5% bacterial reduction in the rinse solution and a > 99.8% reduction on chicken thighs, regardless of the exposure time. Notably, extending exposure time from 1 min to 3 min further enhanced bacterial inactivation, particularly at lower concentrations. These findings underscore the importance of using higher initial PAA concentrations (40–50 mg·L^-1^) and sufficient exposure times (≥ 2 min) to ensure rapid and thorough *Salmonella* inactivation, minimizing cross-contamination risks and enhancing microbial safety in poultry processing systems. In summary, the current study suggests that tracking the changes in both PAA and bacterial levels, not just the initial PAA values, is the key to assessing the efficacy of PAA in inactivating *Salmonella*.

#### 3.6.2. Modelling results

Considering model equations (1) − (3), our objective was to determine the PAA inactivation rate of *Salmonella, k*_*s*_ (L·mg^-1^·min^-1^), on the chicken surface during simulated immersion chilling. The bacterial levels on the chicken, *v* (CFU), is governed by equation (2), which is included here for convenience, where the shed rate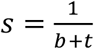 :

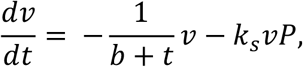

Since in the model, equation (2) decouples from equation (3), we solved for *v* as follows:

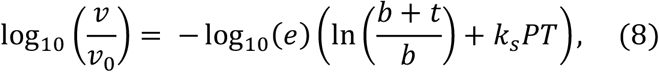

where the term log_10_(*e*) In 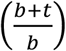 refers to the amount of bacteria shed from the chicken piece into the “chiller water” up to time *T* (min) and the term log_10_(*e*)*k*_*S*_*PT* represents bacteria inactivation via PAA on the chicken surface; recall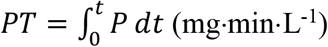. Thus, to estimate the PAA induced bacterial inactivation rate *k*_*s*_ (L·mg^-1^·min^-1^), we used the following equation:

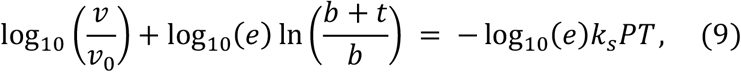

From equation (1) (PAA decay dynamics), for each experimental run (see **section 3.6.1** for details), we used the relationship from the Appendix **Section D** to determine *k*_*TDS*_ as a function of *P*_0_ (set during each experimental run) and hence predict *PT* for respective runs. Furthermore, we set *b* = 0.23 (min) (coming from **Section 3.4.2**), to govern the *Salmonella* shed rate from chicken surface during the chilling simulation and thus estimate the term In 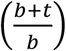 in equation (9). Finally, it should be noted that *v*_0_ (CFU) is the inoculum level on the chicken piece and *v* (CFU) is the resulting bacterial levels measured via the second nBPW rinse during respective runs. Combining this information, a regression analysis was performed using equation (9), which resulted in *k*_*s*_ = 0.13 ± 0.02 (L·mg^-1^·min^-1^) (see **Supplementary Figure 3**). The goodness of fit is illustrated by the residual plot in Appendix **Section C**. A two-sample t-test [MATLAB] revealed that the PAA killing rate on the chicken surface (*k*_*s*_) was significantly different from the PAA killing rate of bacteria in water (*k*_*w*_ = 0.15 ±.02 (L·mg^-1^·min^-1^)). This means, on average, *k*_*s*_ ≈ 0.88 *k*_*w*_, i.e., the surface killing rate is about 70-90% of the PAA killing rate in chiller water without an organic load.

Inputting parameters *k*_*s*_ = 0.13 (L·mg^-1^·min^-1^) (along with *b* = 0.23 (min) and *k*_*w*_ = 0.15 (L·mg^-1^·min^-1^)) into model equations (1) – (3) and normalizing the data in the water by inoculum level (this is the PAA rinse data illustrated in Figures 7 and 8 in section 3.6.1.) to include all experimental runs, we compared the predicted bacterial levels in the water versus the measured levels during the chiller simulations. **Figure 9** illustrates the average model prediction versus average observed bacteria levels (error bars reflect the standard deviation) in the chiller water as a function of PAA contact time, *PT* (mg·min·L^-1^), where the RMSE (across the range of *PT* values) was 0.056 (fraction of inoculum·mL^-1^). The low RMSE value indicates that on average across the range of *PT* values, the model predicts well, even though for larger *PT* values it tends to underpredict killing in the water. Indeed, it may be the case that the Chick-Watson model (equation (5) in **section 3.5.2**) should be modified to capture enhanced inactivation beyond a certain *PT* value. In any case, the results in Figure 9 provide strong evidence for the validity of the determined *average* parameter values of *k*_*s*_, *k*_*w*_, and *b*.

**Fig. 9.**
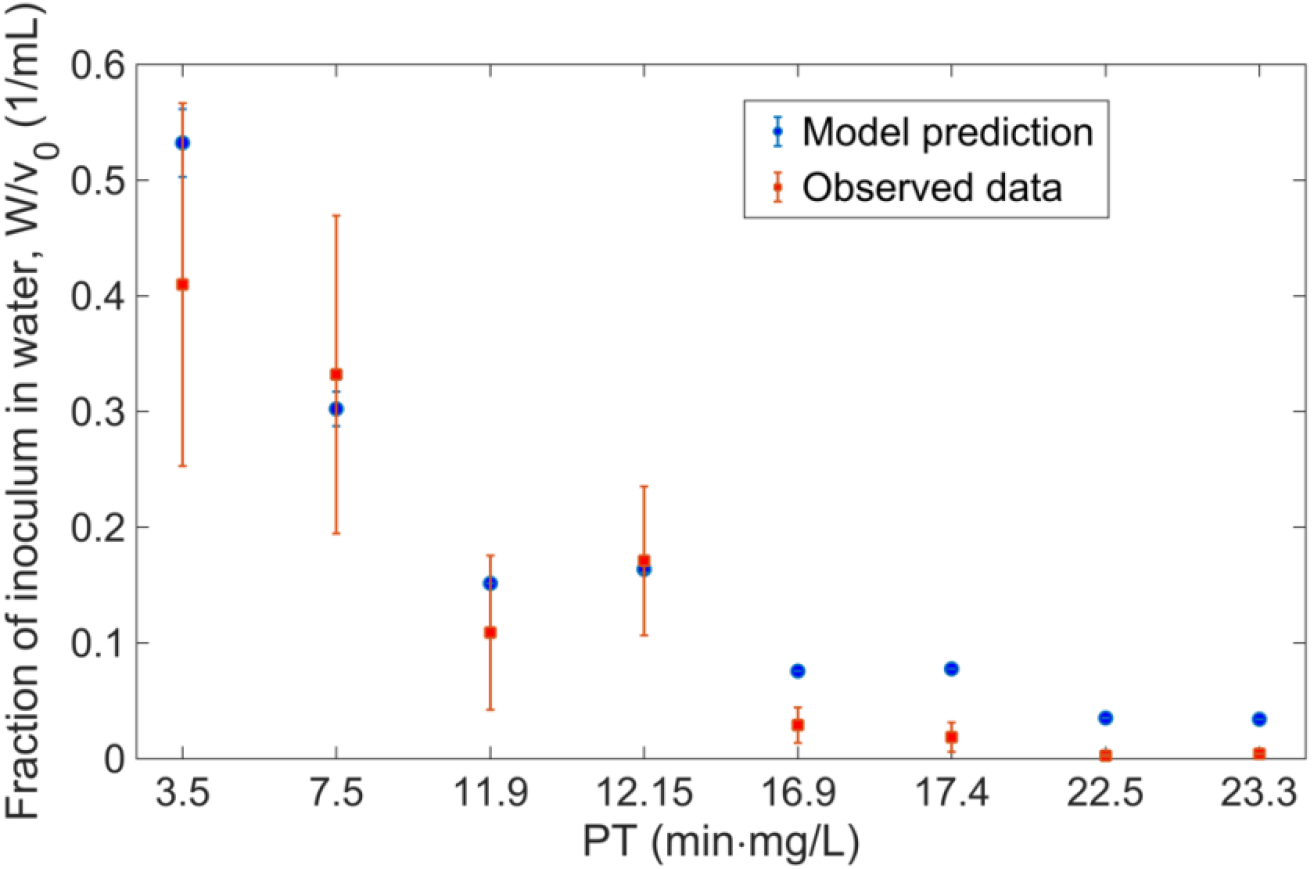
Model prediction of *Salmonella* levels that were shed into the water and survived PAA inactivation

It should be noted that for *PT* values from 3 to 12 (min·mg·L^-1^), there is significantly more variation for observed *Salmonella* levels in the chiller water than for larger *PT* values (Fig. 9). This may be due to differences in PAA tolerance as a function of strain type (a 5-strain cocktail was used), however, the variation in observations as well as the model predictions shrank as *PT* increased. This is logical since beyond a certain PAA concentration or exposure time to PAA, strain differences became negligible and substantial killing of *Salmonella* was observed.

Finally, considering the predictive success of the model shown in Figure 9, we used the model to understand when shedding versus PAA killing on chicken surface dominates the reduction of *Salmonella* on chicken. The experimental time scale for the chilling simulation (data shown in section 3.6.1) was 1 to 3 min. In this context, **Figure 10** shows that on average for PT values < 13 (min·mg·L^-1^), the relative log reduction due to PAA killing was ≲ 25 %, and hence pathogen shedding dominates the reduction of bacterial levels on the poultry. In contrast, for PT values >17 (min·mg·L^-1^), the relative log reduction due to PAA killing was > 60% with PAA inactivation dominating the bacterial reduction. Thus, there is an important transition in PAA efficacy in reducing *Salmonella* on chicken during a simulated chilling process (Fig. 10).

**Fig. 10.**
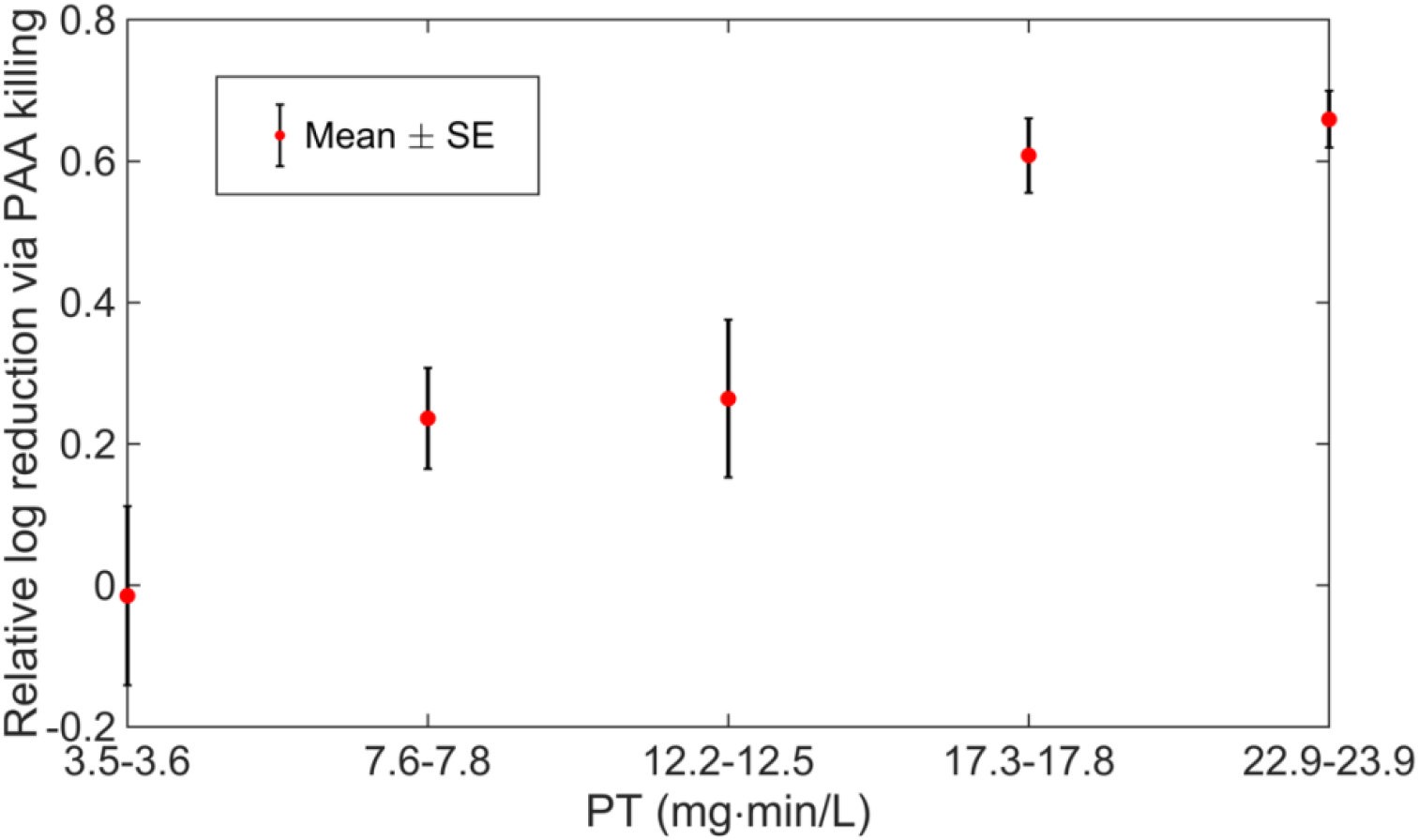
Relative log reduction of *Salmonella* on chicken thighs via PAA exposure. Results are shown as mean value ± standard error. Here total log reduction is defined in terms of both PAA killing and shedding into the chiller water as per the results illustrated in section 3.6.1. Notice that under the experimental conditions (time scale of 1-3 min), PT values from 3-13 (min·mg·L^-1^) correspond to initial PAA levels from 10 to 30 mg·L^-1^ and PT values larger than 17 (min·mg·L^-1^) correspond to initial PAA levels from 40 to 50 mg·L^-1^.

## 4. Conclusions

This study demonstrates the utility of an *experimentally-informed-mechanistic-modeling* approach to describe the mechanisms of bacterial shedding and inactivation during the poultry chilling process. We showed for the first time that the *Salmonella* transfer rate from chicken surfaces to chiller water is a decreasing function of time, with approximately 50% of the total shedding occurring within the first 15 s. This is an important finding as it indicates that the cross-contamination risks associated with *Salmonella* levels in chiller water depend on process time and potentially process time and space, contingent on the type of chiller setup (batch vs counterflow). In addition, without an organic load, we demonstrated that *Salmonella* in water is effectively eliminated by a PAA concentration > 5 mg·L^-1^ with an exposure time ≥ 3 min. In fact, our model results showed that the inactivation rate *k*_*w*_ is independent of bacteria load and is a function of *PT*. The inactivation rate of *Salmonella* on the surface of chicken thighs, *k*_*s*_, during simulated chilling, was verified to be about 70-90% of the killing rate in water with no organic load. Our results also show for the first time that in the post-dissociation of PAA, acetic acid promotes the release of more soluble breakdown products, whereas H_2_O_2_ aggressively oxidizes particulate organic matter.

Interestingly, using our model to compare the critical time scales of *Salmonella* shedding and PAA inactivation both in the water and on chicken surfaces, we found that for lower *PT* values, shedding dominates the removal of *Salmonella* from chicken, but for *PT* values above 17 (min·mg·L^-1^), PAA inactivation plays the dominate role. This result illustrates the synergy of our approach while re-emphasizing the critical role of tracking changes in PAA levels in connection with specific control, i.e., log reduction of *Salmonella* levels. In line with this, our modeling work for PAA levels, validated at the commercial scale, showed that TDS information predicts PAA changes more accurately than COD. While the apparent reaction rate between TDS and PAA, dictated by *k*_*TDS*,_ is reliable under the same context/scale, appropriate calibrations are needed to apply the model across various processing setups. However, the benefit of our work is that TDS can be measured in real-time as opposed to COD, making model tuning quite practical. This study offers an approach and precise measurement of key factors that can influence *Salmonella* behavior during poultry chilling. Understanding these factors is important for making better decisions on how to control pathogens at this crucial stage of poultry processing.

## Appendix

### A. Typical fit result for shedding dynamics

Working from data in section 3.4.1, specifically a whole carcass inoculum level of 3×10^5^ CFU and one minute consecutive carcass rinses (up to 5 times), we illustrate a typical fit result for the shedding model form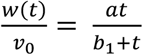, where *v*_0_ = 3×10^5^ (CFU) and *w*(*T*) is the total CFU shed into the water during the time interval [0, *T*]. Results from Levenberg-Marquardt algorithm for nonlinear least squares fitting (**Figure A1**) revealed that for this run, *a* = 0.8577 (fraction of inoculum that can shed into the water) and *b*_1_ = 0.2711 (min), with goodness of fit indicated by *R*^2^ = 0.99 and RMSE = 0.012.

**Figure A1.**
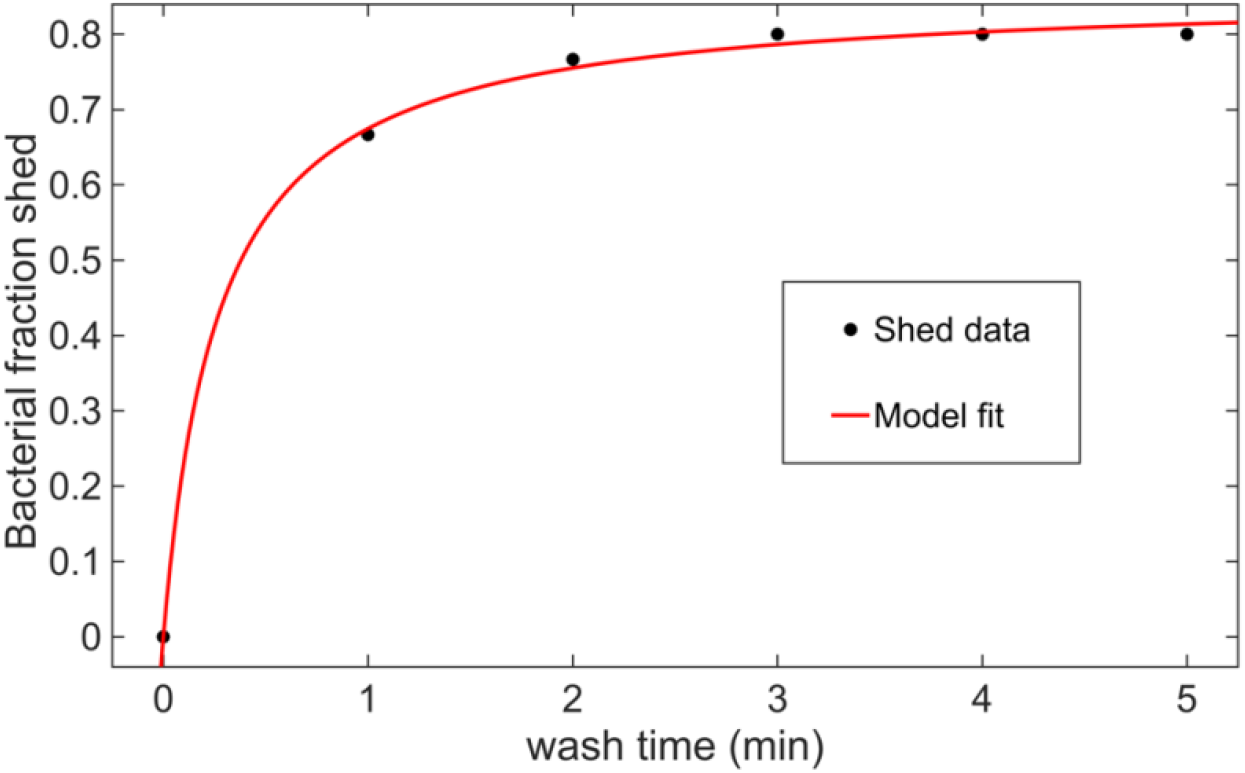
Typical fit result for the model from v_0_ = 3×10^5^ CFU, 1-minute consecutive whole carcass rinses.

The following discussion concerns the treatment of data for experimental runs that included non-detected bacteria levels. Defining *S* to be the total amount of bacteria shed during each experimental run, of the 9 runs for the low initial bacteria loads (average was *v*_0_ = 3×10^5^ CFU), only two from the 1 min rinse duration and one from the 3 min rinse duration had an *S* value that fell outside of 1 standard deviation (4×10^4^ CFU) from the average inoculum level (note that each of these *S* values were slightly less than the average inoculum level). In order to fit the *b* value for these three runs, we considered the following methods: (i) substitute 0 for non-detects, (ii) substitute LOD/2, (iii) substitute LOD/√2, and (iv) multiple imputation (MI) method of taking 100 samples from a uniform distribution on the order of magnitude from 0 to 4, since LOD=10^4^/100 mL [31]. It is worth mentioning that while more sophisticated MI methods relying on maximum likelihood as well as the Kaplan-Meier method have been used in such situations, because of the relatively small number of non-detected data points, our findings below show that the resulting distribution for *b* does not depend on any one of the aforementioned methods [31, 32]. Compared to substituting 0 for the non-detects, replacing the three *b* values from (ii), (iii), or (iv), respectively, shifted the average *b* value (averaged across all experimental runs) by less than 3% and changed upper bounds for the 95% CI by less than 6% and the lower bound by less than 1%. Thus, regardless of how one assigns values to samples below the LOD for these three runs, the effect on *b*, in this context, remains negligible.

### B. Goodness of fit for PAA inactivation rate of *Salmonella* in water (*kw*)

The solution of the Chick-Watson inactivation model (equation (5) in section 3.5.2) was adapted to a suitable form (equation 7 in section 3.5.2) for running regression analysis with PAA inactivation data (coming from section 3.5.1). While Supplementary Figure 2 shows the linear model fit in section 3.5.2., we include here the corresponding residual plot (**Figure B1**). Note that the residuals appear to be randomly distributed about the x-axis, indicating that the appropriateness of a linear model to describe the relationship between PT values and the inactivation ratio.

**Figure B1.**
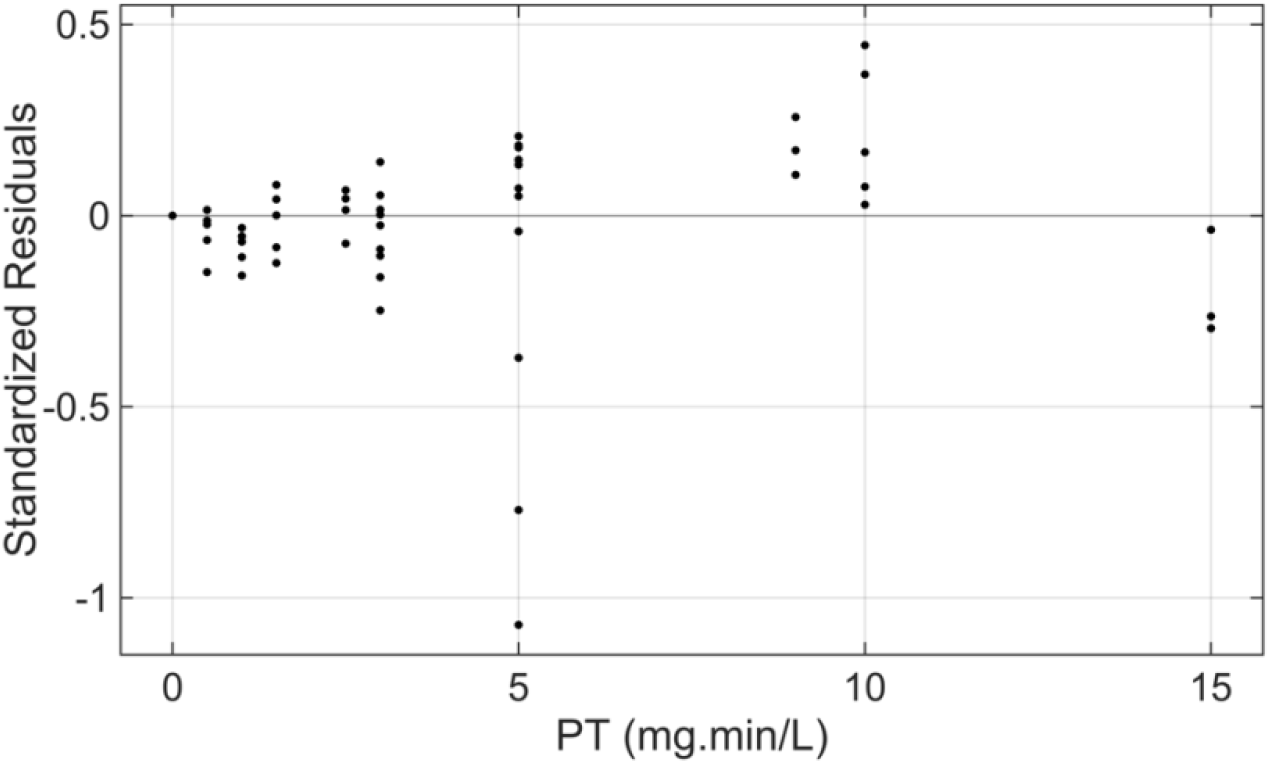
Residual plot for linear inactivation model in equation (7) and Supplementary Figure 2.

In addition, an unbalanced ANOVA analysis was performed in MATLAB to illustrate that the inactivation fraction, 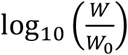, does not depend on the initial PAA concentration *P*_0_ (mg·L^-1^) or the initial inoculum level log_10_ *W*_0_ (CFU·mL^-1^) in the water as the resulting *p*-values were much larger than 0.05 (**Table B1**).

**Table B1.** Results from unbalanced ANOVA analysis.

| Source | SS | Df | MS | F | $p$-value |
| --- | --- | --- | --- | --- | --- |
| $\log_{10} W_0$ | 0.09044 | 1 | 0.09044 | 0.99 | 0.3249 |
| $P_0$ | 0.43557 | 3 | 0.14519 | 1.58 | 0.2034 |
| Error | 5.32226 | 58 | 0.09176 |  |  |
| Total | 5.83435 | 62 |  |  |  |

### C. Goodness of fit for PAA inactivation rate on poultry surface (section 3.6.2)

**Figure C1.**
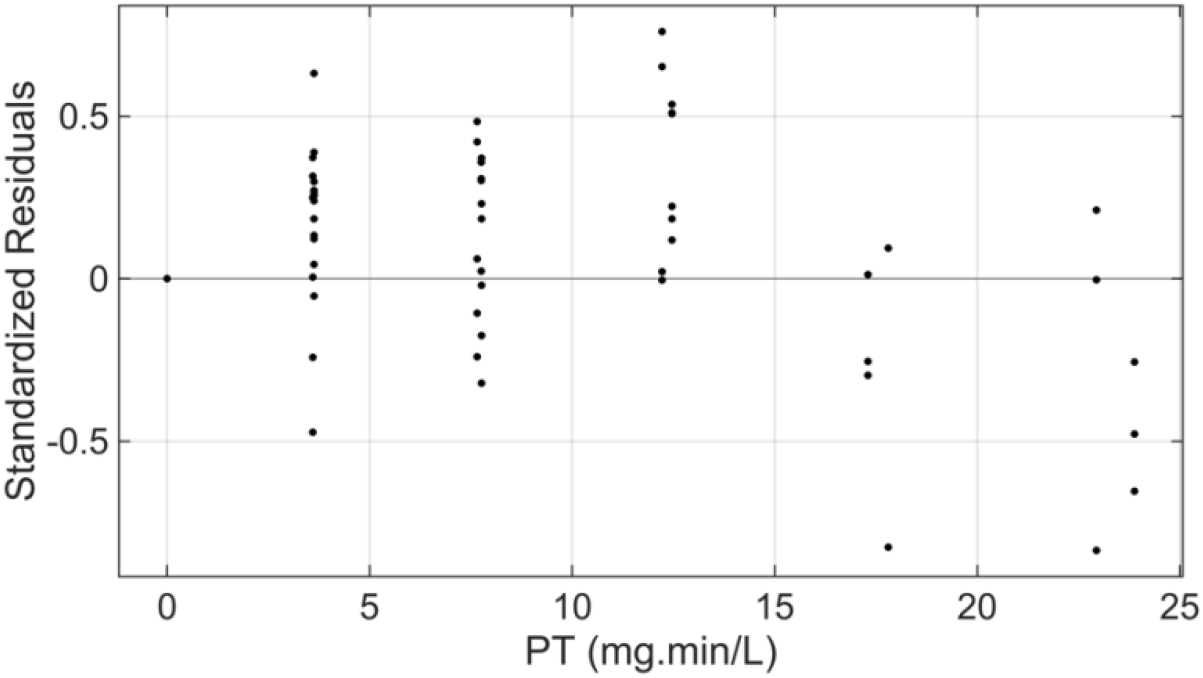
Residual plot corresponding to regression analysis (Supplementary Figure 3) for *Salmonella* inactivation on poultry surface during simulated chilling. Regression results: *R*^2^ = 0.65, RMSE = 0.3459.

### D. PAA decay in 100 mL (100 g thigh piece only; P_0_ = 30, 40, 50, 100, 200)

This section establishes the PAA decay parameter *k*_*TDS*_ as a function of initial PAA concentration (*P*_0_) in 100 ml of DI water in the context of simulating the chilling of a 100 g thigh piece. Here *P*_0_ ranged over 30, 40, 50, 100 and 200 mg·L^-1^ and chilling time lasted at most 3 min. TDS and PAA levels were recorded at *t* = 0, 1, 2 and 3 min, post submerging the thigh piece in the water. Working from equation (1), and setting *X* = TDS level in the tank at time *T*, we have that

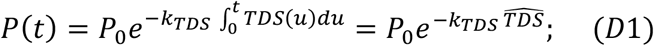

where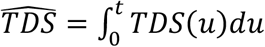. Manipulating equation (D1), we then have:

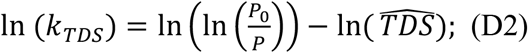

Equation (D2) allows the utilization of all the corresponding data (from experiments in section 3.4) to determine In (*k*_*TDS*_) as a function of *P*_0_ (Figure D1).

**Figure D1.**
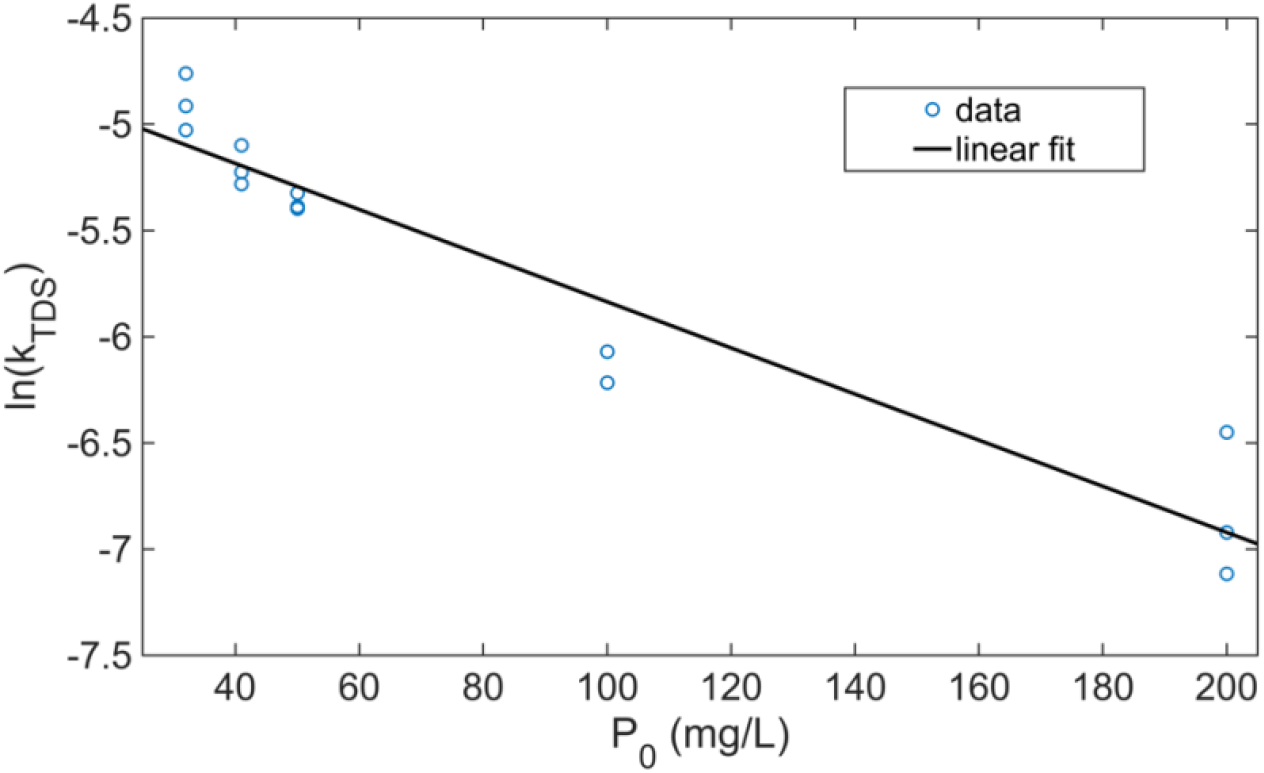
Regression result showing ln (*k*_*TDS*_) as well approximated as a linear function of *P*_0_.

**Figure D2.**
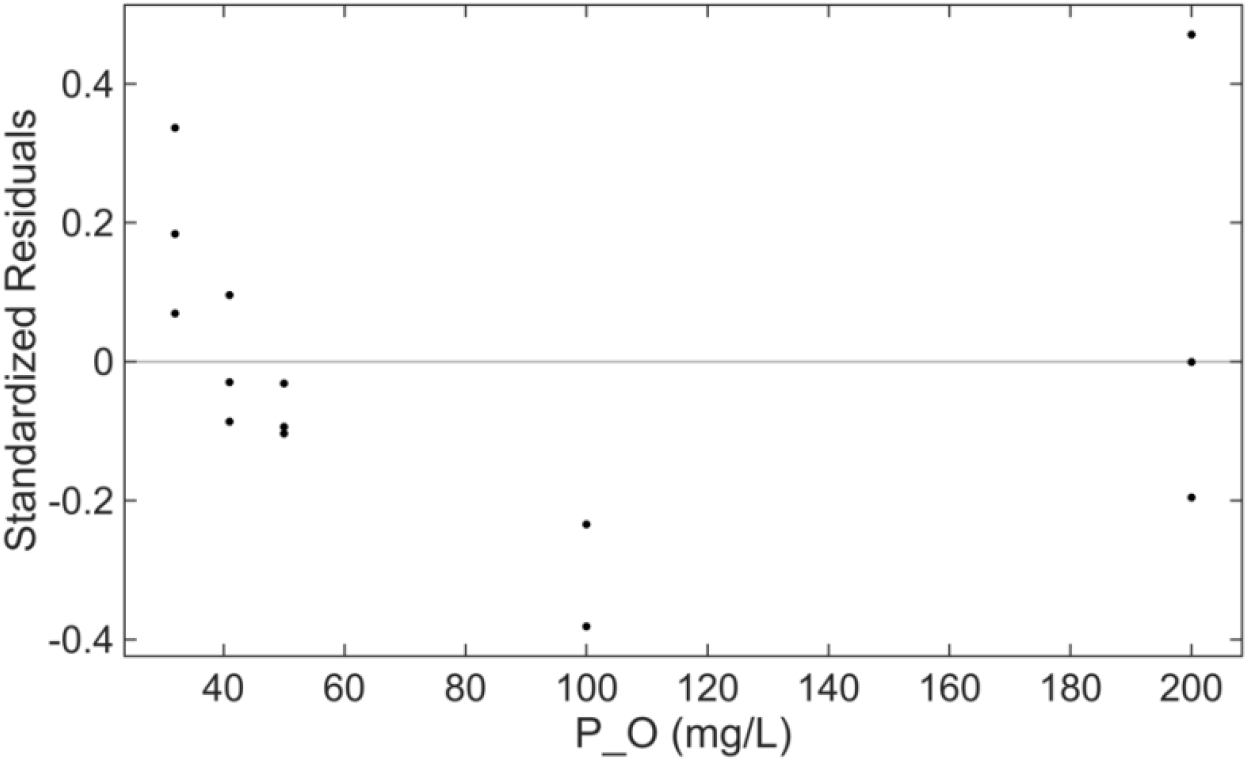
Residual plot for regression analysis in Figure D1.

Figure D1 illustrates the linear model fit

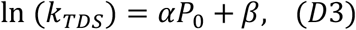

coming from equation (D2), where *α* = −0.011 ± 0.002, *β* = −4.751 ± 0.22 and goodness of fit measures *R*^2^ = 0.91 and RMSE = 0.23. The residual plot (Figure D2) shows that standardized residuals are randomly distributed (Shapiro-Wilk test in MATLAB says normally distributed) about the x-axis, providing justification of a linear relationship between the log of the decay rate and initial PAA concentration (for 30 ≤ *P*_0_ ≤ 200 (mg·L^-1^) and in the context of exudate material coming from 100 mg thigh pieces as measured by TDS levels in the chiller water). Note that we also see this trend with whole carcasses (Table 1 in section 3.1.2.) as *k*_*TDS*_ for *P*_0_ = 200 (mg·L^-1^) is significantly (statistically speaking) less than that for *P*_0_ = 70 (mg·L^-1^). This observation may be due to the lack of skin on one side of the thigh piece as substantially more and most likely different (more reactive with PAA) exudate material from the chicken thigh enters the chiller water than for whole carcasses.

## Acknowledgements

The authors are grateful for the funding support from the National Institute of Food and Agriculture Grant # OHOW-2022-08999, and the Faculty Research and Development Grant from the Office of Research at Cleveland State University. The sponsors have no role in the study design, collection, analysis and interpretation of data, writing of the report, and decision to submit the article for publication. We thank Dr. Jeanine Boulter-Bitzer (OMAFRA, Ontario) for providing chiller water chemistry data from industrial processors. We acknowledge the experimental support from Dr. Mannan Sharma (Environmental Microbial and Food Safety Lab, USDA) for the poultry *Salmonella* serotypes as well as Dr. Elizabeth McMillan and Eric Adams (USDA, Georgia).

## Author Contributions

Vyshnavi Ciluveru: Data curation, Formal analysis, Investigation, Methodology, Validation, Software, Visualization, Writing – original draft, Writing – review and editing.

Jason Simon: Data curation, Formal analysis, Methodology, Software, Validation, Visualization.

Jeffery Farber: Resources, Visualization, Writing – review and editing.

Shawn Ryan: Conceptualization, Formal analysis, Funding acquisition, Methodology, Project administration, Software, Supervision, Visualization, Writing – review and editing.

Chandrasekhar Kothapalli: Conceptualization, Data curation, Formal analysis, Funding acquisition, Investigation, Methodology, Project administration, Resources, Software, Supervision, Validation, Visualization, Writing – original draft, Writing – review and editing.

Daniel Munther: Conceptualization, Data curation, Formal analysis, Funding acquisition, Investigation, Methodology, Project administration, Resources, Software, Supervision, Validation, Visualization, Writing – original draft, Writing – review and editing.

## Notes

### Competing Interest Statement

The authors have declared no competing interest.

