## Supplemental Material for "Peracetic Acid Efficacy and Decay Kinetics in Poultry Processing under Chiller Conditions"

### Supplementary Figures

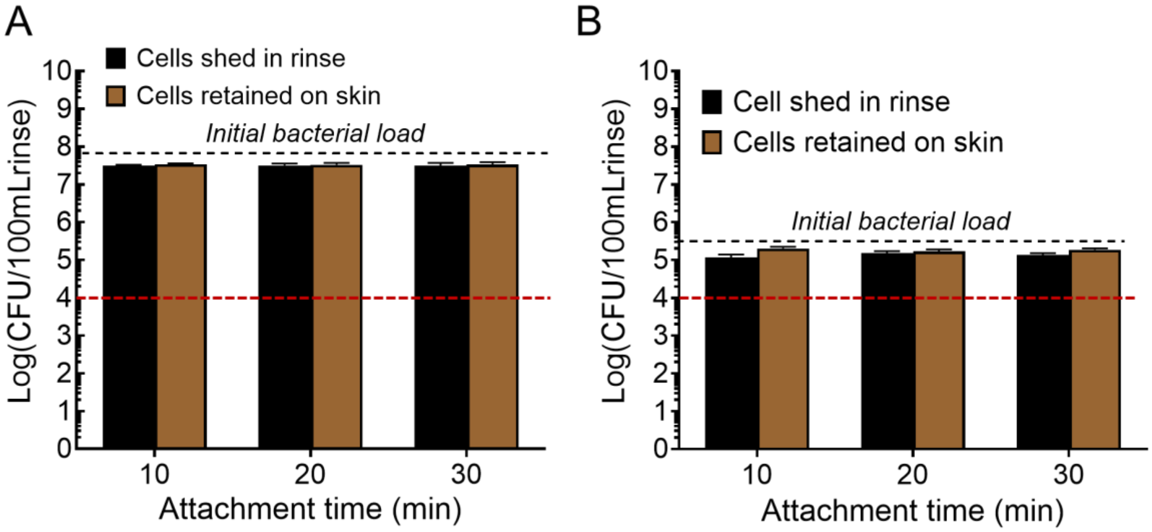


**Supplementary Fig. 1.** *Salmonella* shedding at (**A**) high and (**B**) low load in the absence of PAA. Bacterial shedding and retention are not significantly influenced by attachment duration times (10, 20 or 30 min). The red dotted line represents the limit of detection.

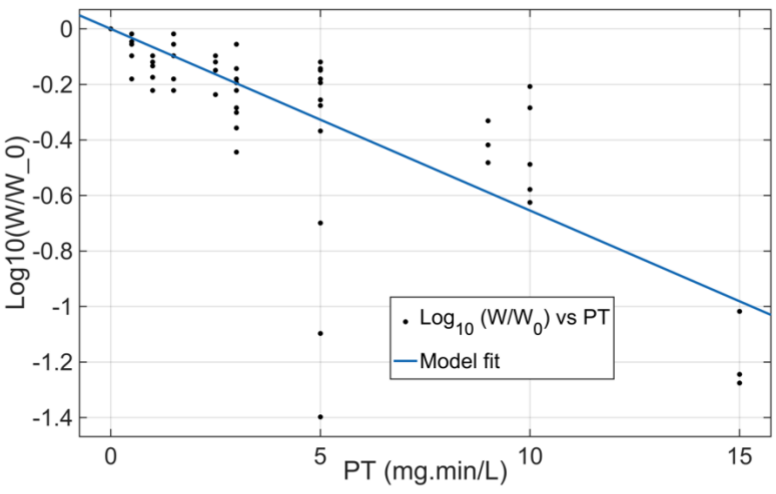


**Supplementary Figure 2.** Regression result for rate of killing in water ($k_w$) with bacteria only via PAA, $R^2$ = 0.58.

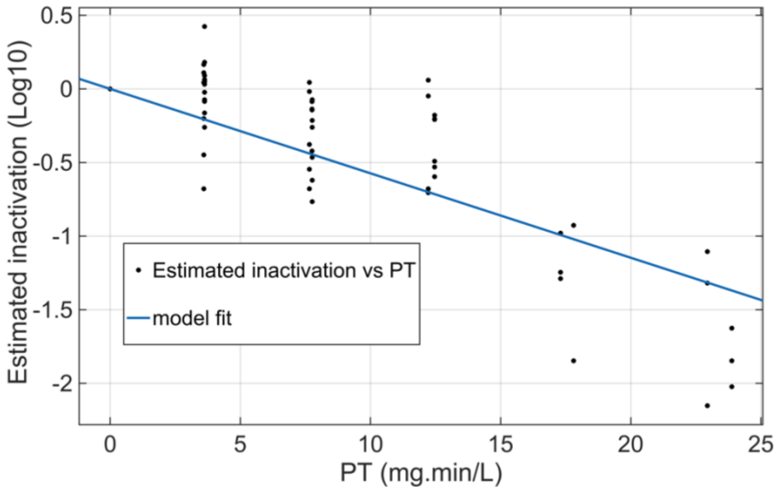


**Supplementary Figure 3**. Model fit result to determine $k_s$; (regression results: $R^2$ = 0.65, RMSE: 0.3459).
